# Structural Categorization and identification of electrostatic interactions in two proposed Human Serum Albumin dimerization patterns and dipyridamole interaction

**DOI:** 10.1101/2025.05.26.656148

**Authors:** Haluk Çetinok, Veyis Karakoç, Erol Erçağ, Yusuf Melih Şekerer, Hasan DeMirci

## Abstract

Human serum albumin (HSA) is a ubiquitous, multifunctional protein responsible for the systemic distribution of both endogenous metabolites and exogenous pharmaceuticals. Its inherent properties, mainly its ability to seep into the tissues and multiple ligand-binding sites, have rendered HSA an attractive vehicle in nanoparticle-based drug delivery systems, particularly in cancer targeting. In this study, we present high-resolution crystallographic data revealing two distinct dimerization patterns of HSA (PDB ID: 9V61), obtained under high-concentration crystallization conditions, in addition to results from dipyridamole dockings. Both dimer types demonstrate extensive interface areas and a significant number of electrostatic interactions. Comparative analysis with previously reported dimer structure (PDB ID: 3JQZ) and other high-interface-area structures, (PDB ID: 5Z0B, PDB ID: 8CKS) indicates similarities in contact regions, but unique residue-level differences in bonding interactions. Interface surface area distribution and space group histograms further support the rarity and potential physiological relevance of the identified dimer forms. Importantly, these dimer configurations do not disrupt Sudlow’s drug-binding sites, which is important as the dipyridamole docking analysis presents strong affinity to Sudlow site I and Sudlow site III, not affecting their utility in engineered drug delivery. Our findings open new avenues for structure-based mutagenesis and nanoparticle design strategies centered on HSA dimerization dynamics.

## 1. Introduction

Human serum albumin (HSA) is an abundantly found globular serum protein produced in the liver which plays a crucial role in transporting metabolites and drug active ingredients throughout the human body [1]. Even though the name may suggest it predominantly localizes intravascularly, HSA is most abundant in extravascular space distributed in tissues and secretions [2]. HSA with a half-life of 19 days approximately completes 28 cycles of dislocating into the tissues and returning to the vascular system via lymphatic vessels [1,2]. This distribution is mainly attributed to the caveolin-dependent Albondin (gp60) mediated transcytosis actively translocating albumin in the capillary lumen to the tissue lumen in order to also transport the nutrients albumin carries into the tissues [3]. In addition to the nutrients, HSA also carries a significant number of exogenous components including the majority of prescribed drugs [4].

Structurally HSA exists as a collection of α-helices (67% α-helices, 10% turns, 23% random coils) exclusively having two main drug binding sites, seven fatty acid binding sites, and two nucleic acid binding sites. [5,6]. Figure 1 shows a scheme summarizing the versatility of HSA highlighting many of its ligands. HSA binding is a dynamic process consisting of the equilibrium between the association and dissociation of ligands thus is dictated by the ligand concentration and free HSA amounts in a specific tissue, for example, as HSA carries the toxic bilirubin to the liver, only 20% of the bilirubin bound to HSA is loaded of in the hepatic capillaries demonstrating how albumin transportation occurs [7].

**Figure 1:**
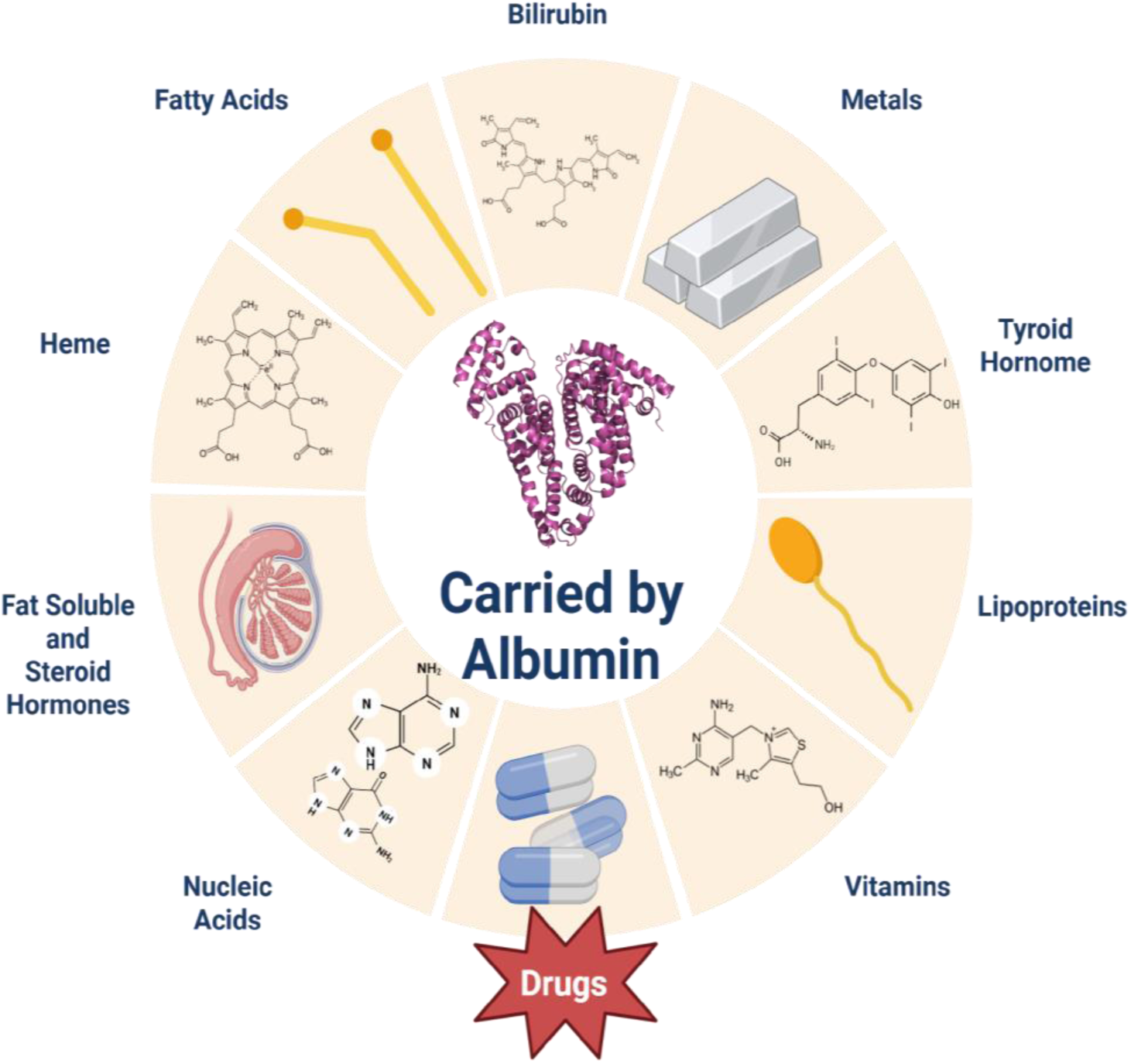
The scheme of some of the metabolites carried by or interacting with HSA. Starting at the top clockwise; Bilirubin, metals, thyroid hormone, lipoproteins, vitamins, drugs, nucleic acids, fat-soluble and steroid hormones, heme, and fatty acids.

As HSA is an important and ample nutrient carrier, multiple types of cancer overexpress the gp60 receptors to better siphon those nutrients rendering HSA an inherently cancer-targeting molecule [8,9]. This disproportionate targeting has given rise to an altogether different area, HSA nanoparticles [10]. HSA nanoparticles (either completely made of HSA or covered with HSA) have at least 2-fold accumulation in tumor tissues as opposed to similar-sized HAS’less nanoparticles [11]. Those nanoparticles can be done via either crosslinking or partially denaturing and renaturing HSA in high concentrations hence HSA self-interactions and multimerization can play an important role in the properties of those nanoparticles [10,11]. HSA consists of three homologous domains each consisting of a double repeating segment kept together via 17 disulfide bridges as presented ın Figure 2. The helices are named depending on their locations regarding those domain repeats.

**Figure 2:**
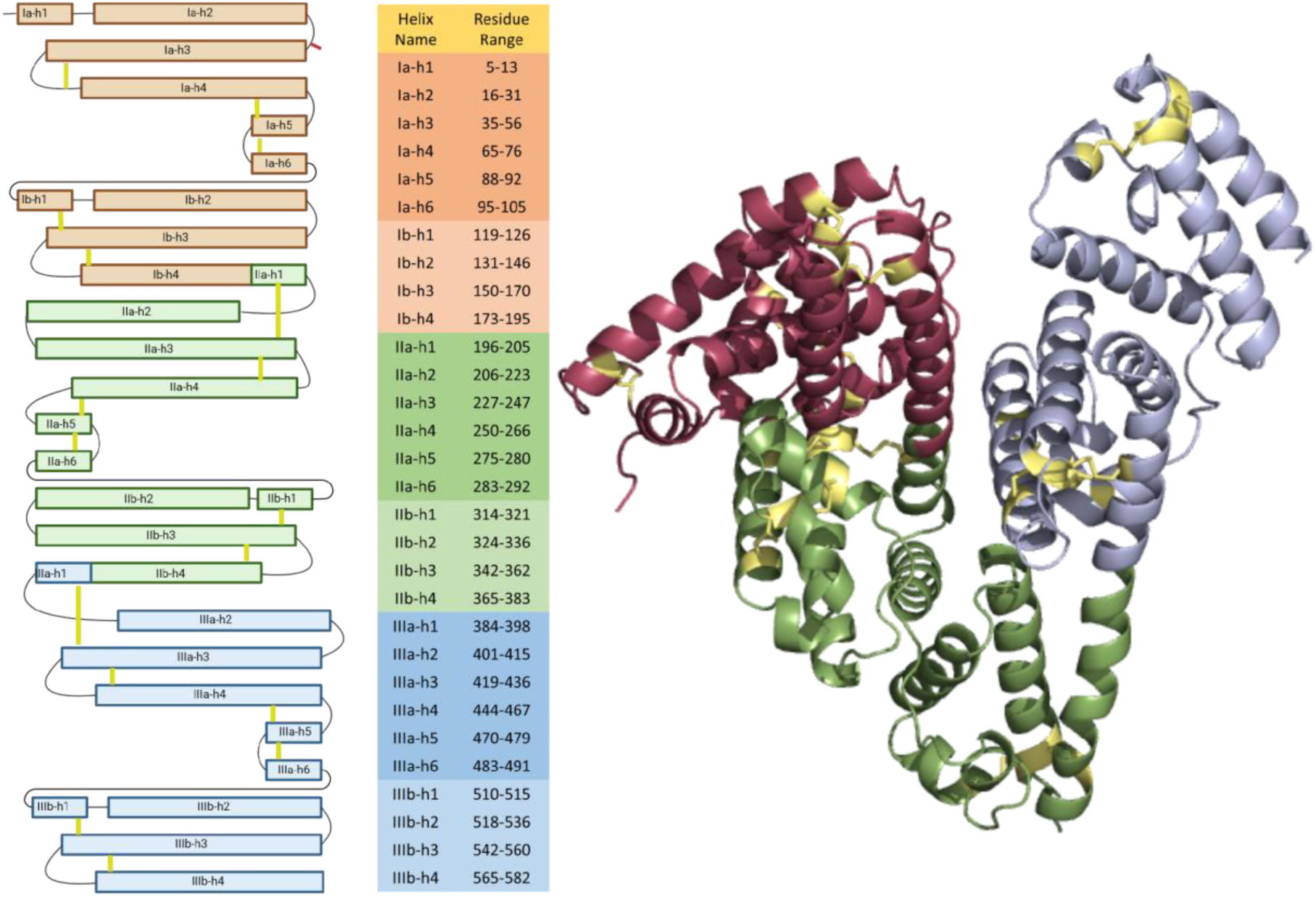
The domains and the helices of human serum albumin with indicated cysteine disulfide bridges are presented. The protein structure presented is the solved structure’s chain A. The subdomains are differentiated via color and the cysteine bridges are placed in their approximate locations.

Both within those repeats and between them there are multiple binding sites for metabolites and drugs [2]. The metabolites and drugs with deposited crystal structures in the Protein Data Bank (PDB) have been aligned to our solved structures Chain A to emphasize the wide distribution of those binding sites on the protein in Figure 3. Fatty acids, drugs, metabolites (like heme and bilirubin), and metals can bind to multiple sites on the protein making every part of its surface relevant for drug transporting and nanoparticle-forming procedures.

**Figure 3:**
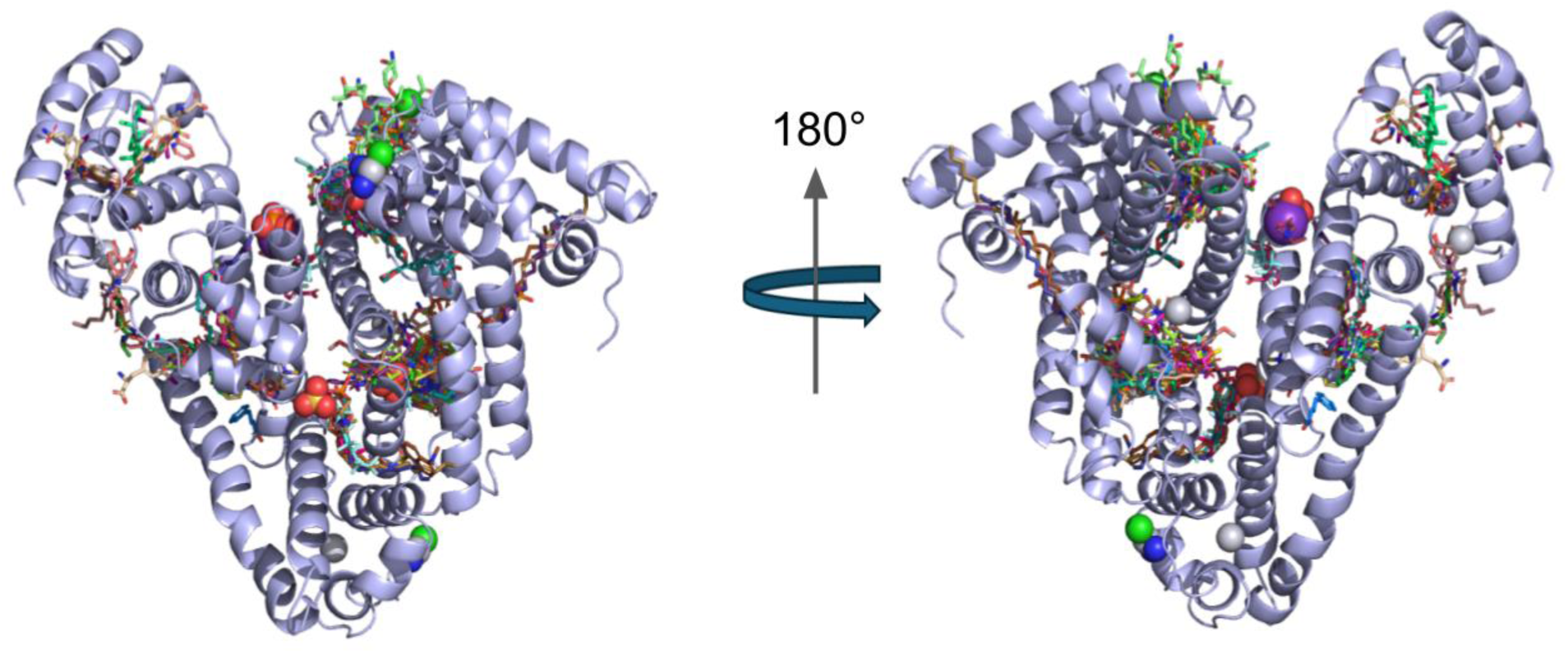
Most known ligands of human serum albumin demonstrated in PDB deposited structures (over 100) aligned to our structure’s chain A emphasizing the versatility and the spread of binding sites on the protein.

HSA can also interact with other proteins as well [12], and in blood, it is at 5% found in a covalent dimeric form with a disulfide bridge between two HSA monomers [13,14]. Even though there is little evidence of HSA forming reversible dimeric in physiologic conditions [13,14], recombinantly dimerized HSA and HSA multimerized in nanoparticles have higher drug targeting efficiency [11,15] hence artificial and/or in vivo dimerization patterns may be utilized for improving nanoparticle efficiency and play a role in mutagenesis targeting. In crystallization, the dimers occurring can either have biological significance or be a crystallization artifact, usually, the area of the interface between the monomers of the dimer and the interaction strength differentiates between them [16]. Non-biological dimer interfaces tend to be smaller (<1000 Å) with little electrostatic interactions. The only other established HSA reversible dimer (PDB accession code: 3JQZ) has a similar surface area to our interfaces (1718.9Å to 1695.1Å) and comparable electrostatic interactions (28 to 24) [17]. While usually the driving force between protein dimers is hydrophobic interactions, the interaction being forced by electrostatic interaction, especially on higher concentrations, is not unheard of [18]. As HSA dimerisations and self-interactions can influence both nanoparticle dynamics and targeting properties, discovering possible dimerization patterns could introduce new parameters to both fields as mutagenesis targets and may offer insights into new conditions in which the HSA nanoparticles could be generated.

Human serum albumin (HSA) contains several binding regions, as shown in Figure 1, with different properties that make this protein a significant factor for drug distribution in the bloodstream, as several pharmaceutical drugs tend to bind to specific regions of the protein. For instance, drugs such as warfarin or phenylbutazone have a higher affinity towards Sudlow Site I. On the other hand, diazepam, ibuprofen, or naproxen demonstrate a higher affinity towards Sudlow site II [19] [20]. In the present study, molecular docking simulations of dipyridamole have been performed at the Sudlow Site I of the IIA subdomain of HSA.

The drug active ingredient dipyridamole is mainly prescribed as an antiplatelet agent as one of its functions is inhibiting phosphodiesterases leading to the accumulation of signaling molecules cAMPs and cGMPs hence inhibiting the platelet activity [21]. Even though dipyridamole is not prescribed other than an antiplatelet agent, its other effects include increasing unassisted patency of synthetic arteriovenous hemodialysis grafts [22], inhibiting mengovirus DNA replication [23], and increased cytotoxic activity against cancer cells compared to normal cells [24] making it a drug repurposing candidate. For a drug’s behavior within the patient, half-life, and distribution, and aforementioned affinities, mainly HSA’s tendency to accumulate in cancer tissues, the drug interactions with the serum proteins can be crucial in manipulating said properties as well as developing novel therapeutic delivery methods regarding those drugs [25].

In this research, the initial aim was to observe the real-space binding of the HSA protein with the drug dipyridamole. However, in the experimentation that binding was not observed and instead, a potential homodimerization pattern for HSA was discovered and discussed, and circling back to dipyridamole dockings were conducted and compared with other established bindings.

## 2. Materials and methods

The HSA protein was acquired commercially and then purified using gel filtration. The purified protein was then incubated with the drug dipyridamole and put through microdrop crystallization in terasaki plates. The acquired crystals were further analyzed in x-ray diffraction using our home source Turkish DeLight and analyzed using the software crysalispro and Phenix. Once it became clear that the drug dipyridamole was not a part of the crystal docking analyses were conducted.

### 2.1 Purification of HSA

The HSA used was commercially available intravenously administered medicine Albuman by Centurion brand (200mg/mL). The Protein was then filtered via gel filtration using superdex200 size exclusion chromatography as a buffer exchange step with the buffer consisting of 150mM NaCl and 20mM Tris-HCl Ph: 7.35. Using Amicon concentrators the pure protein was concentrated up to 99 mg/mL,, this final concentration was confirmed using a nanodrop spectrophotometer. The protein was then added to the drug dipyridamole until its solubility limit (with the undissolved drug visible) and stored in a 4°C fridge. Figure 4 gives a simplified summary of the HSA purification and crystallization.

**Figure 4:**
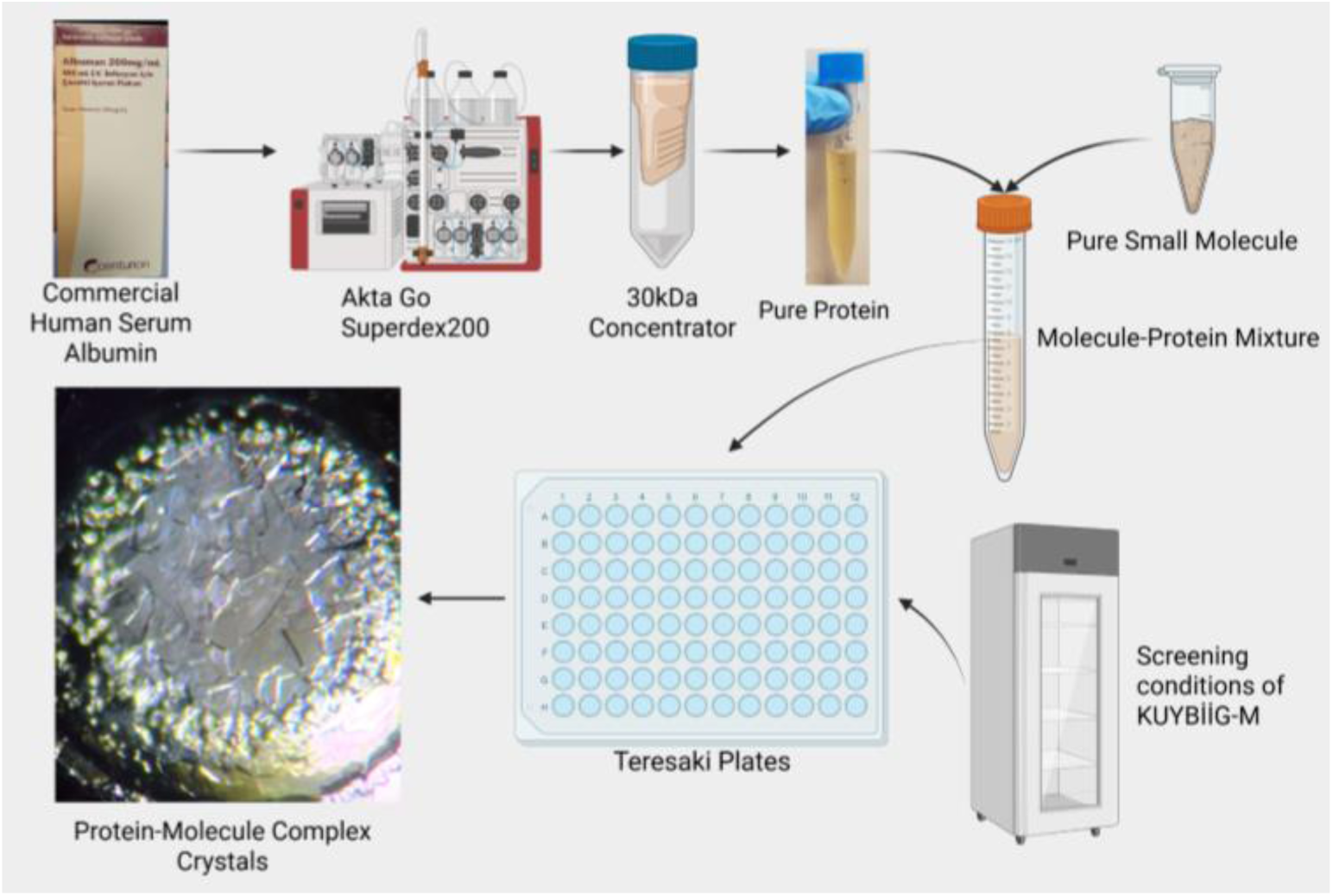
Flowchart of the crystallization process

### 2.2 Crystallisation of HSA

The protein sample was mixed with over 3000 crystallization solutions in the Kuybiig-M (Koç University Structural Biology and Innovative Drug Design Center) library in equal volume (0.83µL to 0.83µL) and covered with a 16.6µL of paraffin oil. Among all those the protein crystallised in commercial crystallization solution crystal structure #39 consisting of 2.0M Ammonium Phosphate Monobasic in 0.1M Tris 8.5 pH solution. The crystals were first observed after three weeks of room temperature incubation in the terasaki plate in a dark environment.

### 2.3 Data collection

Ambient temperature X-ray crystallographic data was collected using Rigaku’s XtaLAB Synergy R Flow XRD system according to the Turkish DeLight X-ray source protocols [26]. Multiple crystals were screened using the modified adapter of the XtalCheck-S plate reader. During the data collection due to protein precipitation, the crystals were not visible yet upon searching in the well multiple areas gave x-ray diffraction data. The duration of exposure time was optimized to minimize the potential radiation damage caused by X-rays. Diffraction data were collected on different days for a total of approximately 10 hours. The detector distance was set at 100.0 mm, and the scan width and the angle of the plate were arranged to cover around 80°.

### 2.4 Data processing

The diffraction data were set up in CrysAlisPro to complete the automated data reduction. The collected data was then merged using the profit merge process with CrysAlisPro 1.171.42.59a software to produce an integrated reflection dataset (*.mtz) file for further analysis [26].

The Phenix phaser was used for molecular replacement and refinement to fully position the folded protein chains into the three-dimensional electron map of the .mtz file. The resulting .pdb protein structure file was then manually refined in Coot version 0.9 and visualized in PyMol software.

### 2.5 Pisa analyses

The bond lengths, counts, and interface surface areas were calculated and generated using PDBePISA online software. All deposited PDB unique HSA structures (excluding non-crystal structures and ones with non-HSA peptides) were put through the software one by one for the generation of the statistics presented in Figure 5 and Figure 6.

**Figure 5:**
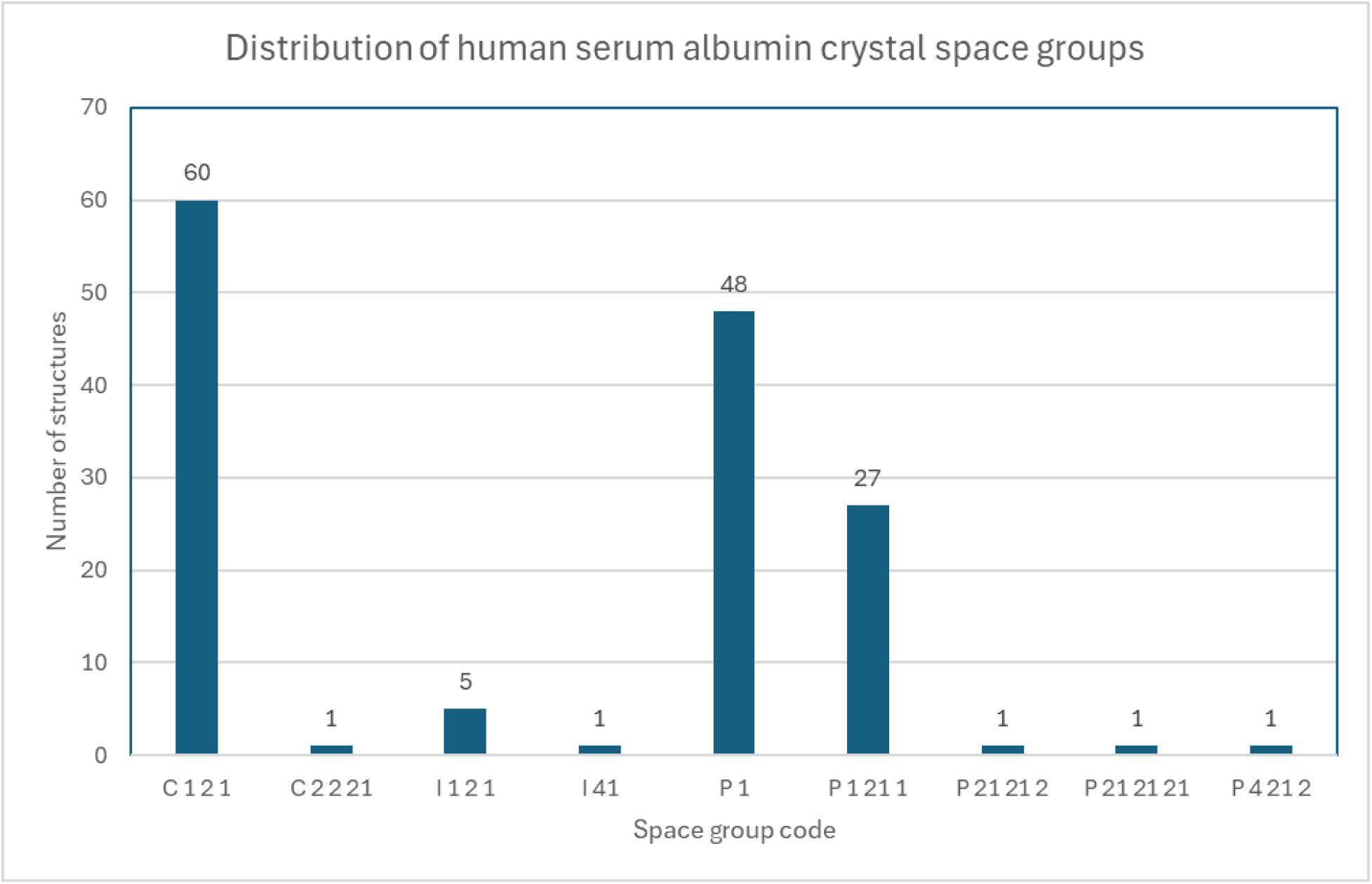
The histogram depicting the distribution of the space groups of human serum albumin crystals in all crystal structures in PDB, heterogeneous (having more than one unique peptide chain) and non-crystal (cryo-EM and such) are excluded. The two proposed dimers in this paper (A-B and C-C) are included and counted in the I 1 2 1 space group.

**Figure 6:**
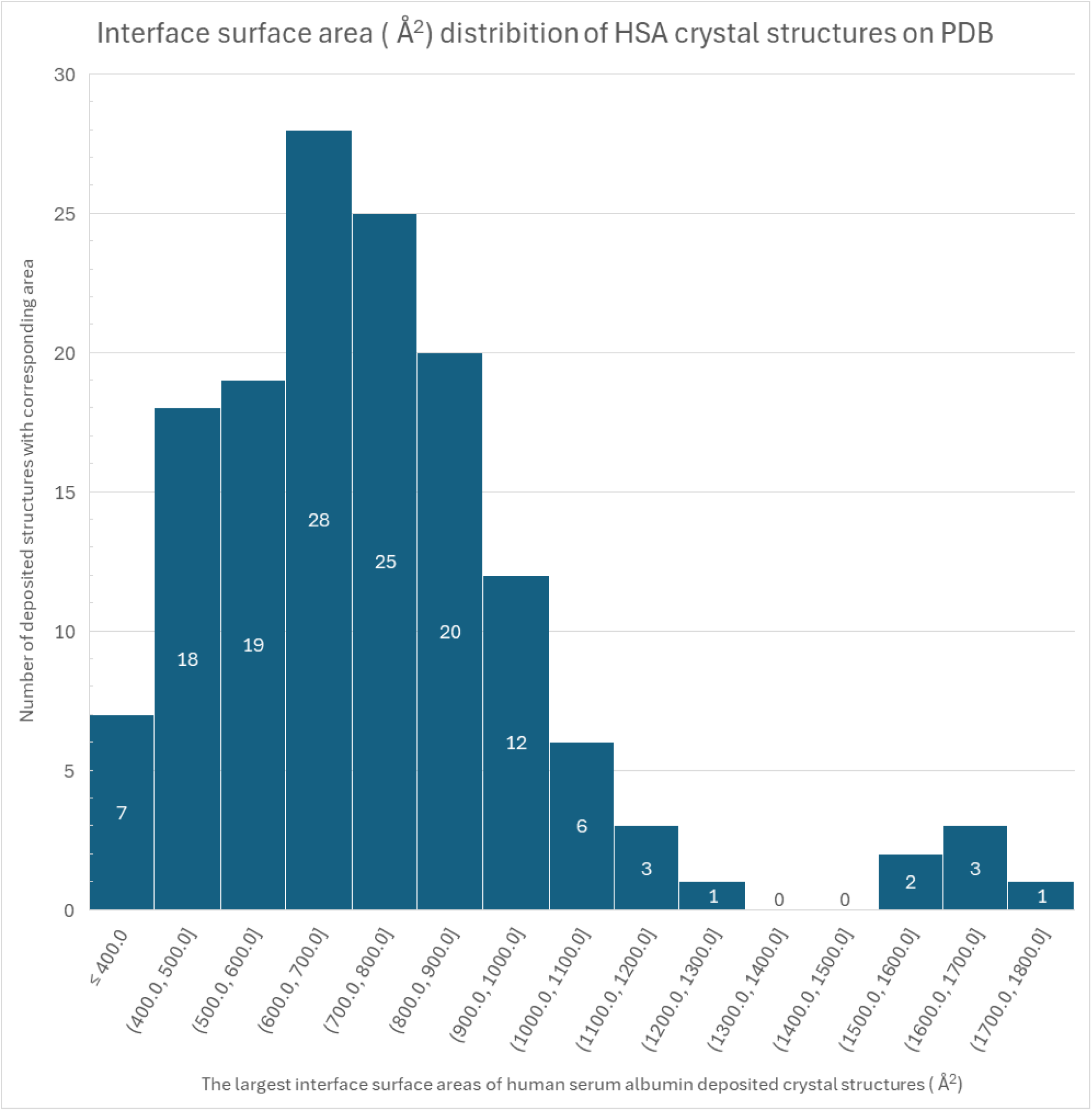
The histogram depicting the distribution of the interface surface areas between human serum albumin chains in all hsa crystal structures in PDB, heterogeneous (having more than one unique peptide chain) and non-crystal (cryo-EM and such) are excluded. The two proposed dimers in this paper (A-B and C-C) are included and counted in the 1600.0-1700.0 Å^2^ range.

### 2.6 Figure and Table Generations

The figures were generated using Bio Render software and the tables were made using Microsoft Excel software. The Figures depicting protein foldings were prepared using Pymol visualization software.

### 2.7 Molecular Docking

Molecular docking calculations were performed using Autodock Vina software by Scripps Research Institute [27]. Three crystal structures of HSA with ligands bound to different binding sites have been downloaded from the Protein Data Bank (PDB). For the Sudlow I binding site, Sudlow II binding site, and Sudlow III binding site, 1H9Z (HSA protein complexed with warfarin) [28], 2BXH (HSA protein complexed with indoxyl sulfate) [19], and 4L8U (HSA protein complexed with 9-amino-camptothecin) had been downloaded, respectively. All of the crystal structures have resolutions higher than 2.50Å.

The proteins that have been downloaded have been prepared using MGL tools. Firstly, the water molecules in the crystal structure have been deleted, secondly, the amino acid residues with missing hydrogen atoms have been protonated. Finally, the Kollman charges have been added for the proteins.

Following the protein preparation, the ligands in the crystal structures were docked into the binding sites in order to validate the docking processes by calculating the root mean squared deviations (RMSD) by comparing the positions of identical atoms between the crystalized ligand and the docked ligand.

Finally, the dipyridamole molecule was modeled and minimized by the MMFF94s forcefield in Avogadro software [29] and prepared in Autodock. After ligand preparation, dipyridamole was docked into the active site that had been designated previously. Discovery Studio Visualizer [30] software was used to analyze the docking results.

## 3. Results

Other than a trimer [31] and another dimer (PDB accession codes: 5Z0B and 8CKS) there is no other crystal structure among the hundreds of structures in PDB with an interaction surface larger than 1300 Å different from our two proposed dimers. In addition to the interaction surface area, our structures (PDB ID: 9V61) also have out-of-ordinary space groups in their crystals, Our solved structures and 8CKS are *I 1 2 1* and the only reversible dimer 3JQZ is the only *I 41*. How rare having higher than 1300Å^2^ interface area occurrences and the given particular space groups are presented in Figure 5 and Figure 6 respectively.

The histogram of the interface areas given in Figure 8 has an empty range between 1300Å^2^ and 1500Å^2^ giving the visual of two separate normal distributions.

It can be understood from Figure 7 that the space-filling of the repeating unit has two distinct dimerization patterns, the first being between chains A and B colored blue and green in Figure 7.b and the other being where two asymmetric units interlock into each other between their chain C’s. The alignment of the Chains of the structure is presented in Figure 8.

**Figure 7:**
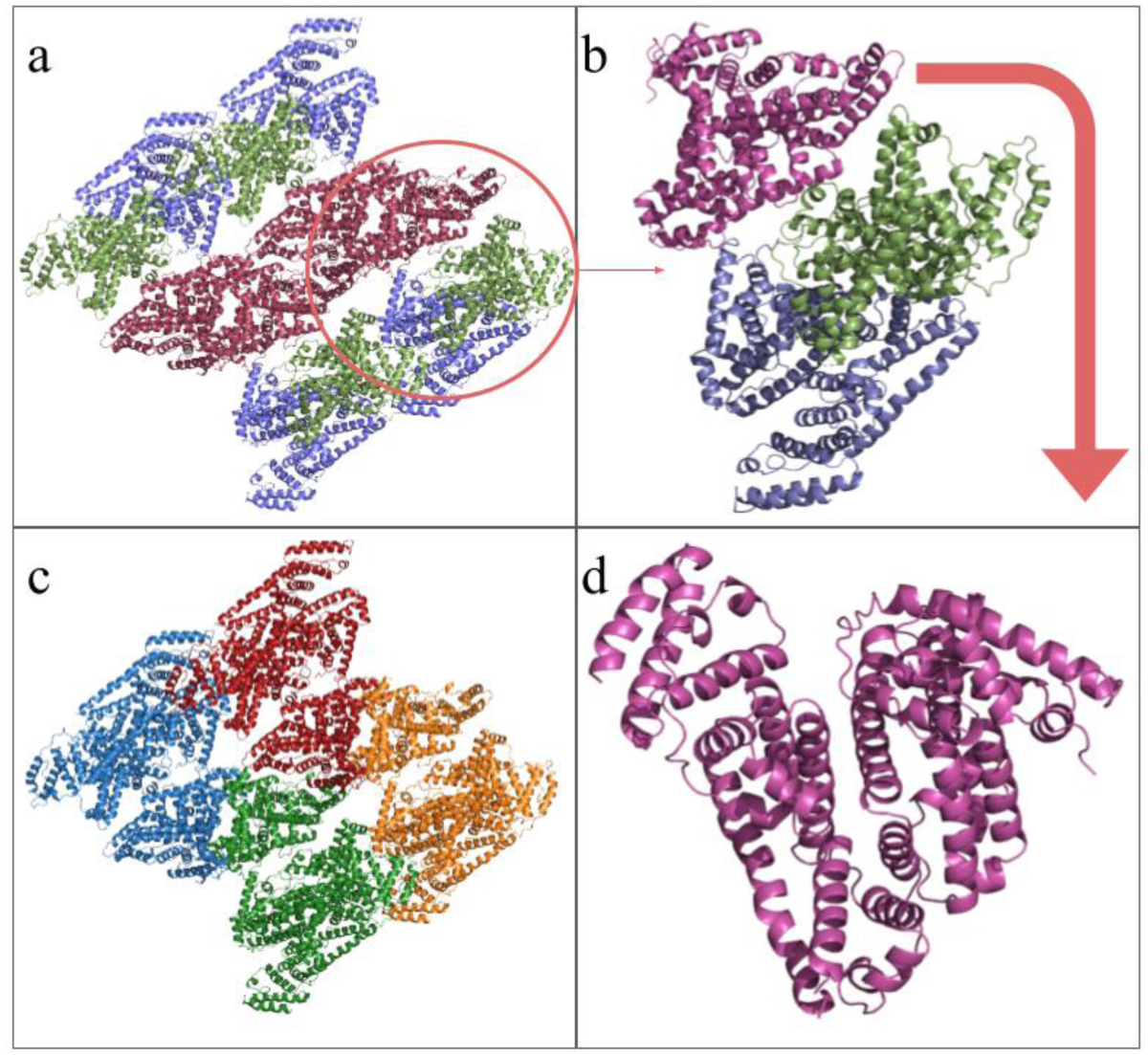
The repeating and asymmetric units’ components are colored and demonstrated as a flowchart. (a) The repeating unit of the solved structures consists of 4 asymmetric units each containing three HSA proteins. Colored by chains in the asymmetric unit (b) A Single asymmetric unit consisting of three individual protein chains is extracted from the repeating unit. (c) The repeating unit with individual asymmetric units is colored differently. (d) Single HSA protein (Chain C) extracted from the asymmetric unit.

**Figure 8:**
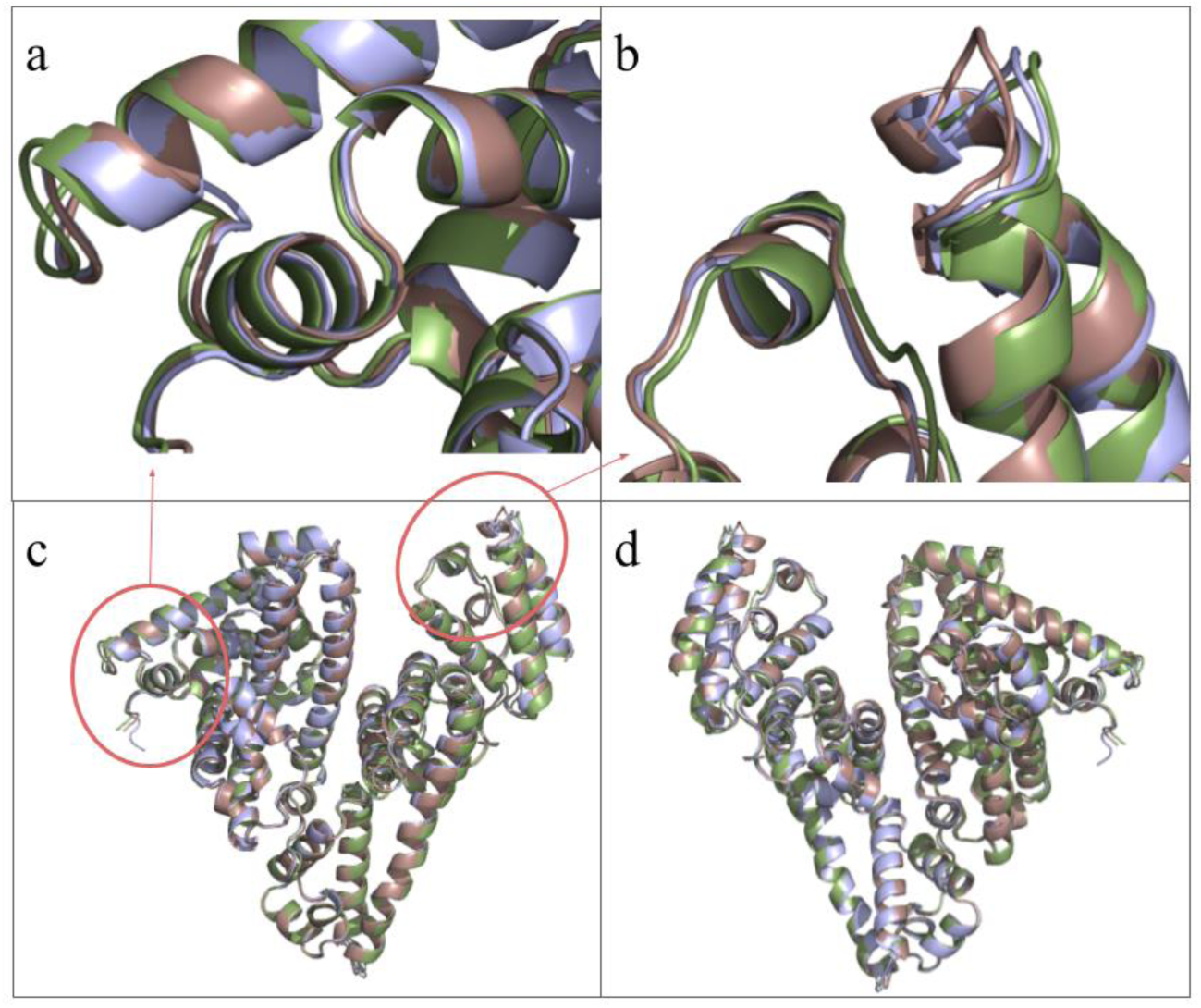
Three chains of our molecule aligned at the residues 365-398. Chain is colored green B and chain C is colored brown. (a) This panel focuses on the slight shift between the helices of Ia-H3 and Ia-H4. (b) This panel focuses on the slight shift between the helices of IIIb-H3 and IIIb-H4. (c) This panel shows the overall alignment from the front. (d) overall alignment from back

The solved structure (PDB ID: 9V61) is presented in Figure 7. How the asymmetric unit assembles to the repeating unit is presented as a flowchart, the entire repeating unit filling the *I 1 2 1* space is presented in Figure 7. a with all unique protein chains colored individually. The single asymmetric unit is extracted into Figure 7.b. The repeating unit consists of four asymmetric units as seen in Figure 7.c with each asymmetric unit consisting of three protein chains colored differently. A single HSA chain is shown in Figure 7.d.

Figure 8. a and Figure 8.b focus on slight differences between the unique chains in the asymmetric unit while Figure 8.c and Figure 8.d offer front and back views of the total alignment structure. RMSD values are A-B 0.461, A-C 0.522, and B-C 0.625.

The overall structure of these dimers in relation to each other is presented in Figure 9, and individual independent visualizations of our proposed dimers are presented in Figure 10.

**Figure 9:**
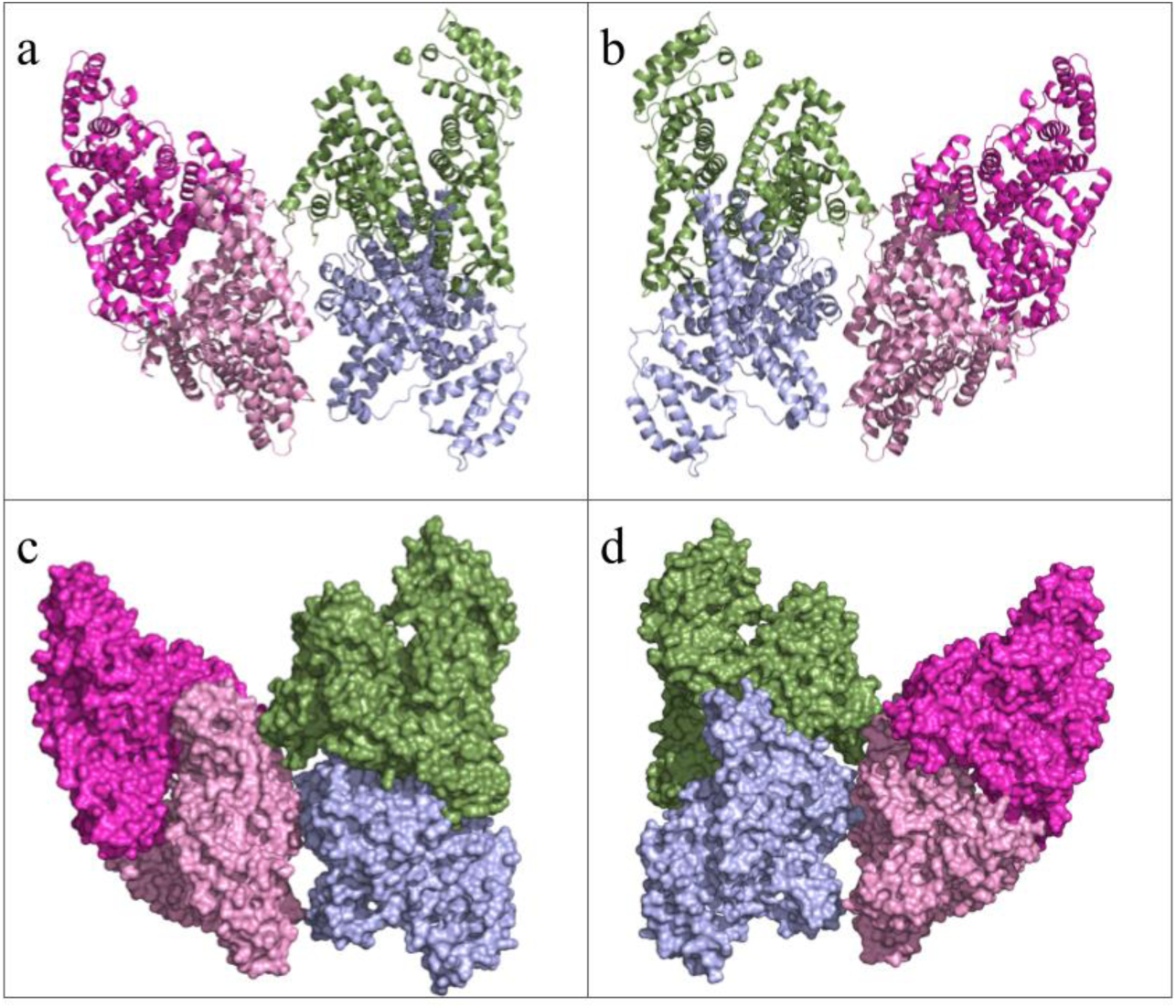
The two dimerized structures solved. Chain A is colorized blue, Chain B is colorized green, Chain C is colorized light pink and the other asymmetric unit’s chain C is colorised in magenta. (a) Four albumins are arranged in two distinct dimer formations in cartoon representation, front view. (b) Four albumins are arranged in two distinct dimer formations in cartoon representation, back view. (c) Four albumins are arranged in two distinct dimer formations in surface representation, front view. (d) Four albumins are arranged in two distinct dimer formations in surface representation, back view.

**Figure 10:**
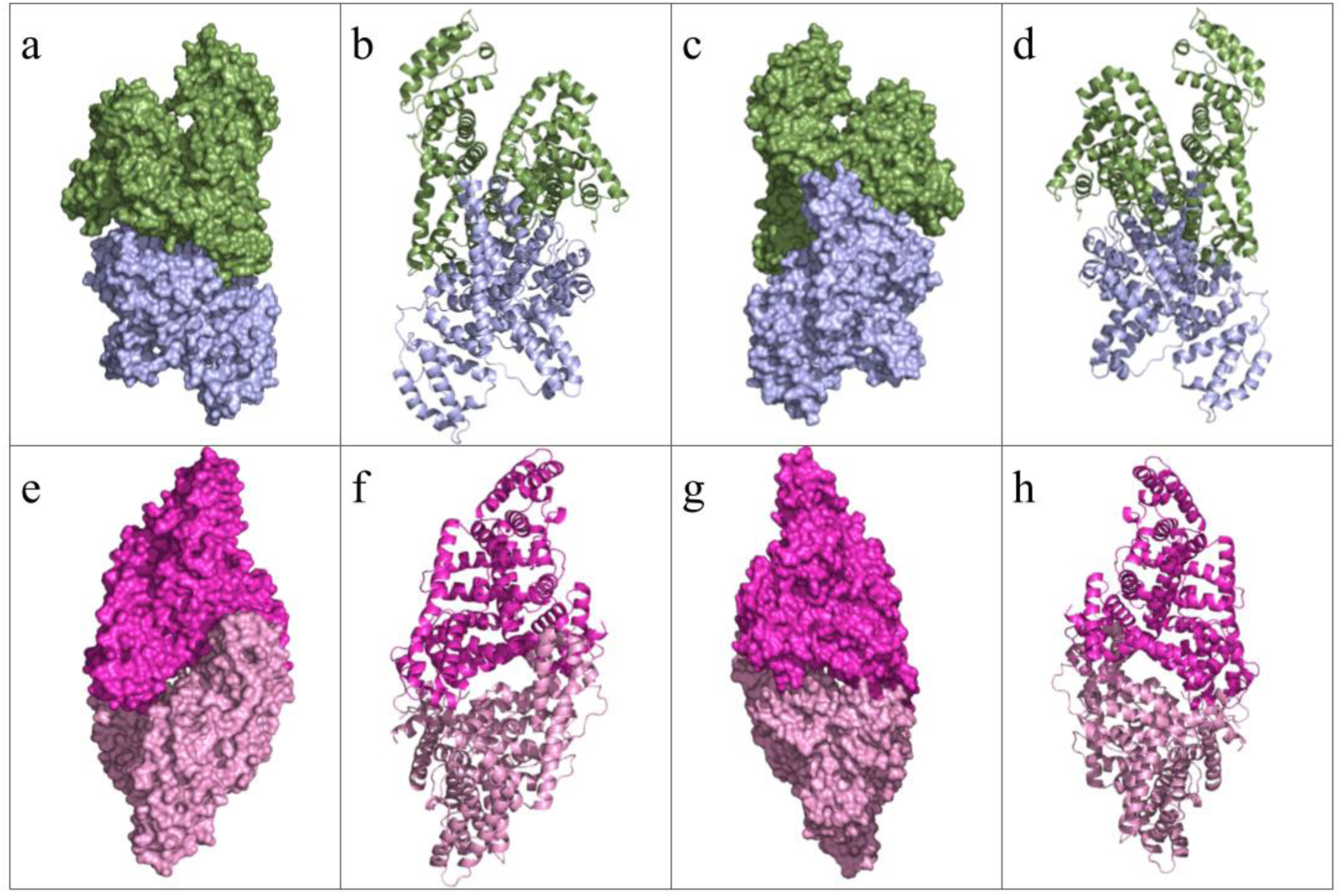
The two dimerized structures solved. Chain A is colorized blue, Chain B is colorized green, Chain C is colorized light pink and the other asymmetric unit’s chain C is colorised in magenta. (a) The dimer of the chains A and B, shown in the front view as surfaces. (b) The dimer of the chains A and B, are shown in front view as cartoons. (a) The dimer of the chains A and B, shown in the back view as surfaces. (b) The dimer of the chains A and B, are shown in the back view as cartoons. (e) The dimer of the chains C1 and C2, shown in the front view as surfaces. (f) The dimer of the chains C1 and C2, shown in the front view as cartoons. (g) The dimer of the chains C1 and C2, shown in the back view as surfaces. (h) The dimer of the chains C1 and C2, shown in the back view as cartoons.

The two distinct dimerization patterns, A-B and C1-C2 were aligned having the RMSD value of 0.716. The salt bridge contributors of both A-B and C1-C2 dimers were presented in Table 1 as a list and as visuals in Figure 11 and Figure 12. The hydrogen bonds between A-B are listed in Table 2 and shown in Figure 13, the C-C dimer hydrogen bonds are listed in Table 3 and visualized in Figure 14. The solved structures (PDB ID: 9V61) proposed dimers’ alignment to the mentioned similar structures are presented in Figure 15, with RMSD values being 3JQZ: 3.784, 5Z0B: 1.0013 and 8CKS: 2.976

**Figure 11:**
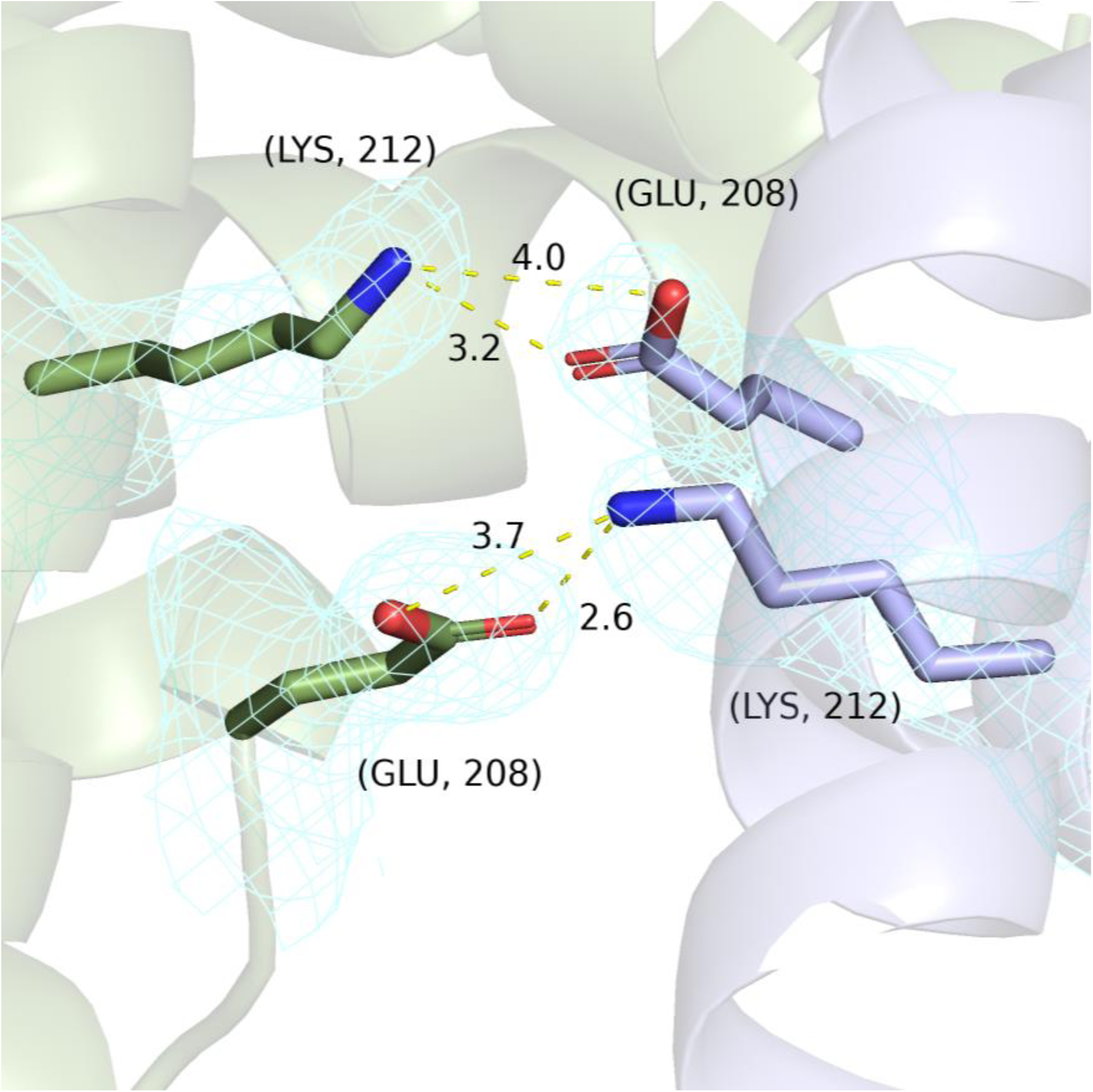
The salt bridges presented by PISA between chains A and B are shown with labels and distances. The cyan mesh corresponds to the electron cloud, the green-backboned protein is Chain B and the blue-colored protein is Chain A.

**Figure 12:**
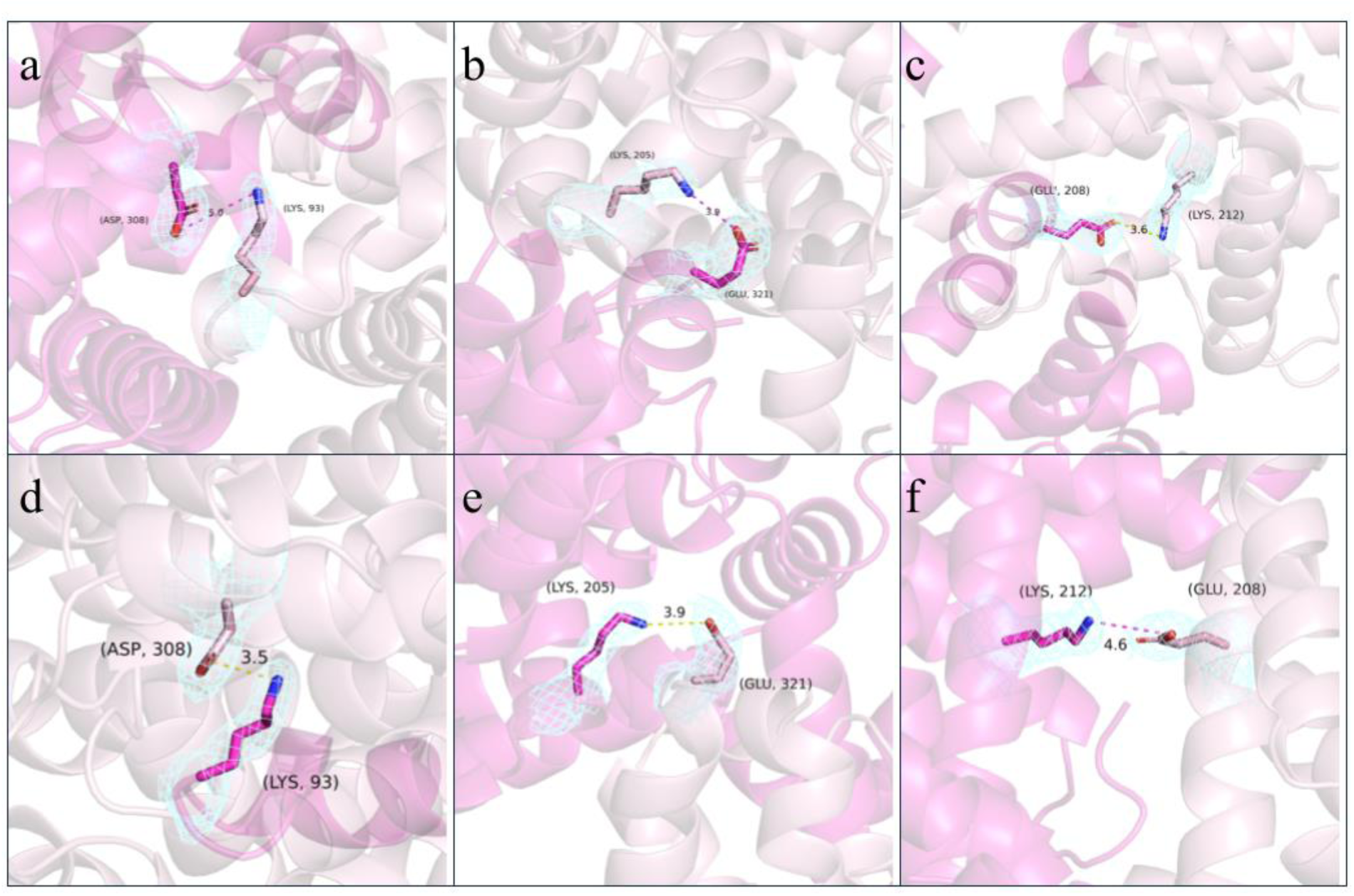
The salt bridges presented by PISA between two chain C’s are shown with labels and distances. (a) The salt bridge numbered 1 (Chain C1 Lys93NZ and Chain C2 Asp308OD1) is shown. (b) The salt bridge numbered 2 (Chain C1 Lys205NZ and Chain C2 Glu321OE2) is shown. (c) The salt bridge numbered 3 (Chain C1 Lys212NZ and Chain C2 Glu208OE1) is shown. (d) The salt bridge numbered 4 (Chain C1 Asp308OD1 and Chain C2 Lys93NZ) is shown. (e) The salt bridge numbered 5 (Chain C1 32Glu321OE2 and Chain C2 Lys205NZ) is shown. (f) The salt bridge numbered 6 (Chain C1 Glu208OE1 and Lys212NZ) is shown.

**Figure 13:**
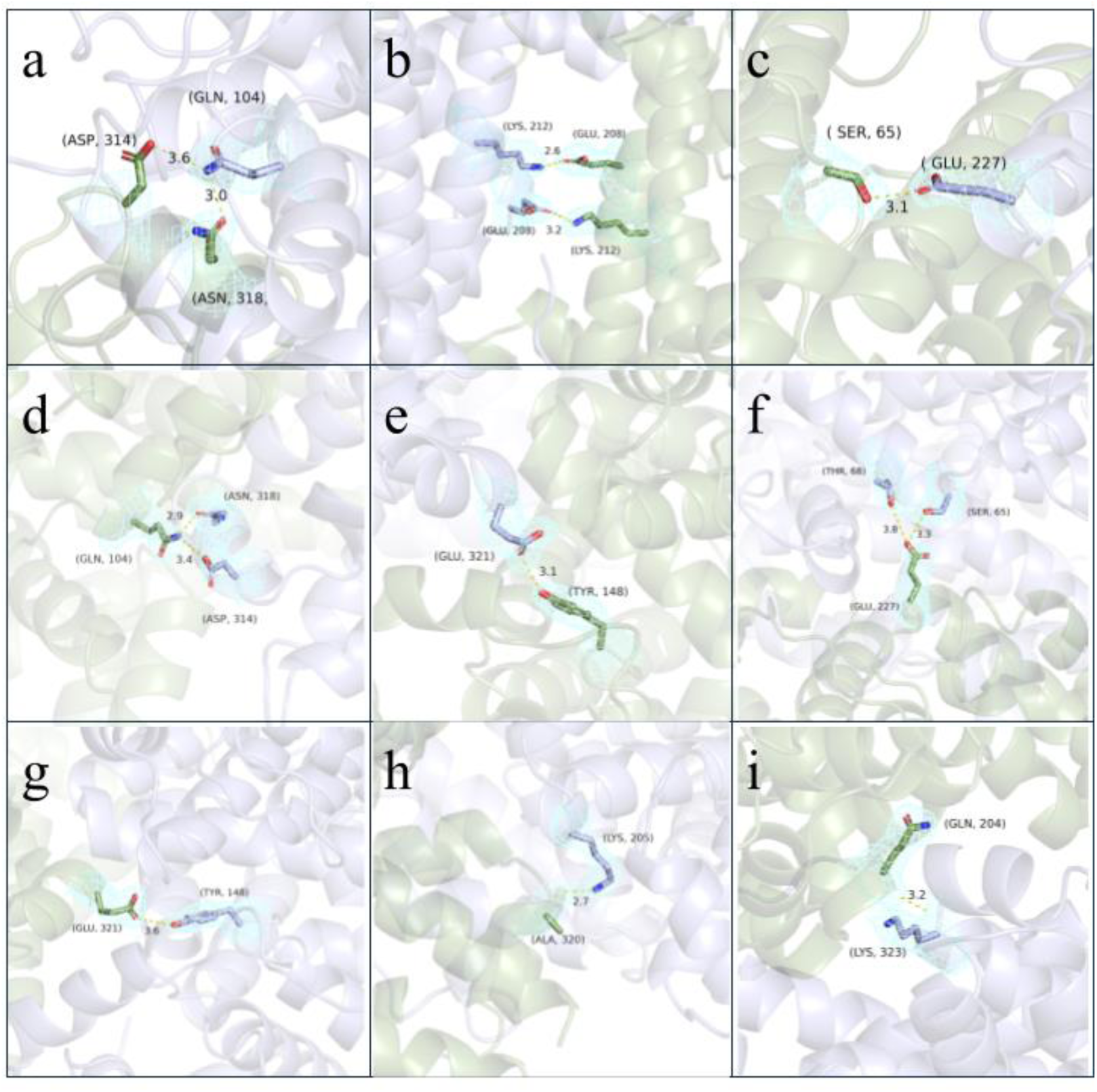
The hydrogen bonds presented by PISA between chains A and B are shown with labels and distances. The blue mesh is the observed electron cloud, prepared in Coot software. (a) The hydrogen bonds numbered 10 (Chain B Asn318OD1 and Chain A Gln104NE2) and 11 (Chain B Asp314OD2 and Chain A Gln104NE2) are shown. (b) The hydrogen bonds numbered 2 (Chain B Lys212NZ and Chain A Glu208OE1) and 14 (Chain B Glu208OE1 and Chain A Lys212NZ) are shown. (c) The hydrogen bond numbered 3 (Chain B Ser65OG and Chain A Glu227OE1) is shown. (d) The hydrogen bonds numbered 4 (Chain B Gln104NE2 and Chain A Asp314OD2) and 5 (Chain B Gln104NE2 and Chain A Asn318OD1) are shown. (e) The hydrogen bond numbered 6 ( Chain B Tyr148OH and Chain A Glu321OE1) is shown. (f) The hydrogen bonds numbered 8 (Chain B Glu227OE2 and Chain A Ser65OG) and 9 (Chain B Glu227OE2 and Chain A Thr68OG1) are shown. (g) The hydrogen bond numbered 12 (Chain B Glu321OE1 and Chain A Gln104NE2) is shown. (h) The hydrogen bond numbered 13 (Chain B Ala320O and Chain A Lys205NZ) is shown. (i) The hydrogen bond numbered 15 (Chain B Gln204O and Chain A Lys323N) is shown.

**Figure 14:**
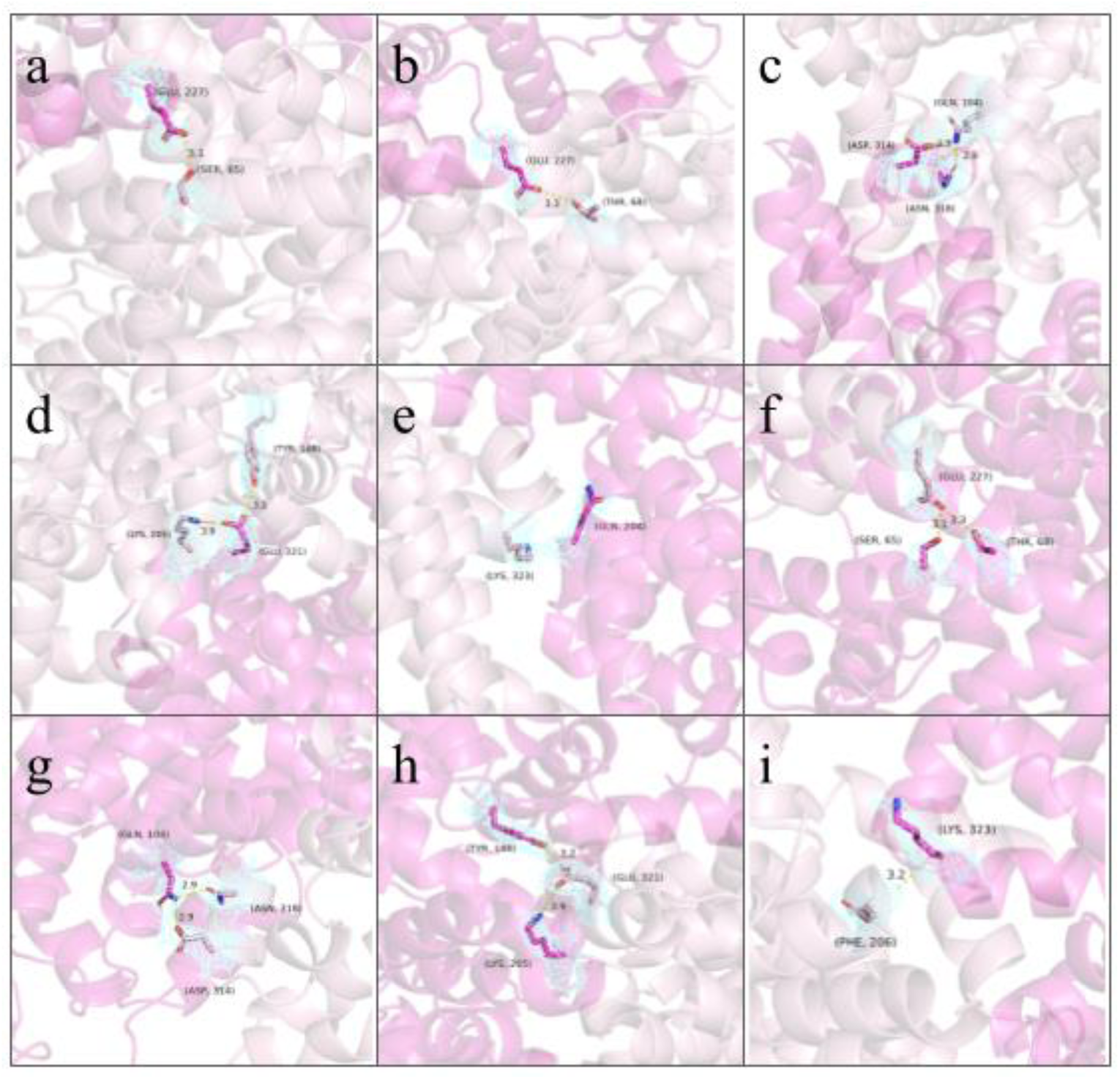
The hydrogen bonds presented by PISA between two chain C’s are shown with labels and distances. The blue mesh is the observed electron cloud, prepared in Coot software. (a) The hydrogen bonds numbered 2 ( Chain C1 Ser65OG and Chain C2 Glu227OE1) and 3 (Chain C1 Ser65OG and Chain C2 Glu227OE2) are shown. (b) The hydrogen bond numbered 4 (Chain C1 Thr68OG1 and Chain C2 Glu227OE2) is shown. (c) The hydrogen bonds numbered 5 (Chain C1 Gln104NE2 and CHain C2 Asn318OD1) and 6 (Chain C1 Gln104NE2 and Chain C2 Asp314OD2) are shown. (d) The hydrogen bonds numbered 7 (Chain C1 Tyr148OH and Chain C2 Glu321OE1) and 8 (Chain C1 Lys205NZ and Chain C1 Gln204O) are shown. (e) The hydrogen bond numbered 9 (Chain C1 Lys323N and Chain C2 Gln204O) is shown. (f) The hydrogen bonds numbered 11 (Chain C1 Glu227OE1 and Chain C2 Ser65OG), 12 (Chain C1 Glu227OE2 and Chain C2 Ser65OG) and 13 (Chain C1 Glu227OE2 and Chain C2 Thr68OG1) are shown. (g) The hydrogen bonds numbered 14 (Chain C1 Asp314OD2 and Chain C2 Gln104NE2) and 15 (Chain C1 Asn218OD1 and Chain C2 Gln104NE2) are shown. (h) The hydrogen bonds numbered 16 (Chain C1 Glu321OE1 and Chain C2 Tyr148OH) and 17 (Chain C1 Glu321OE2 and Chain C2 Lys205NZ) are shown. (i) The hydrogen bond numbered 18 (Chain C1 Gln204O and Chain C2 Lys323N) is shown.

**Figure 15:**
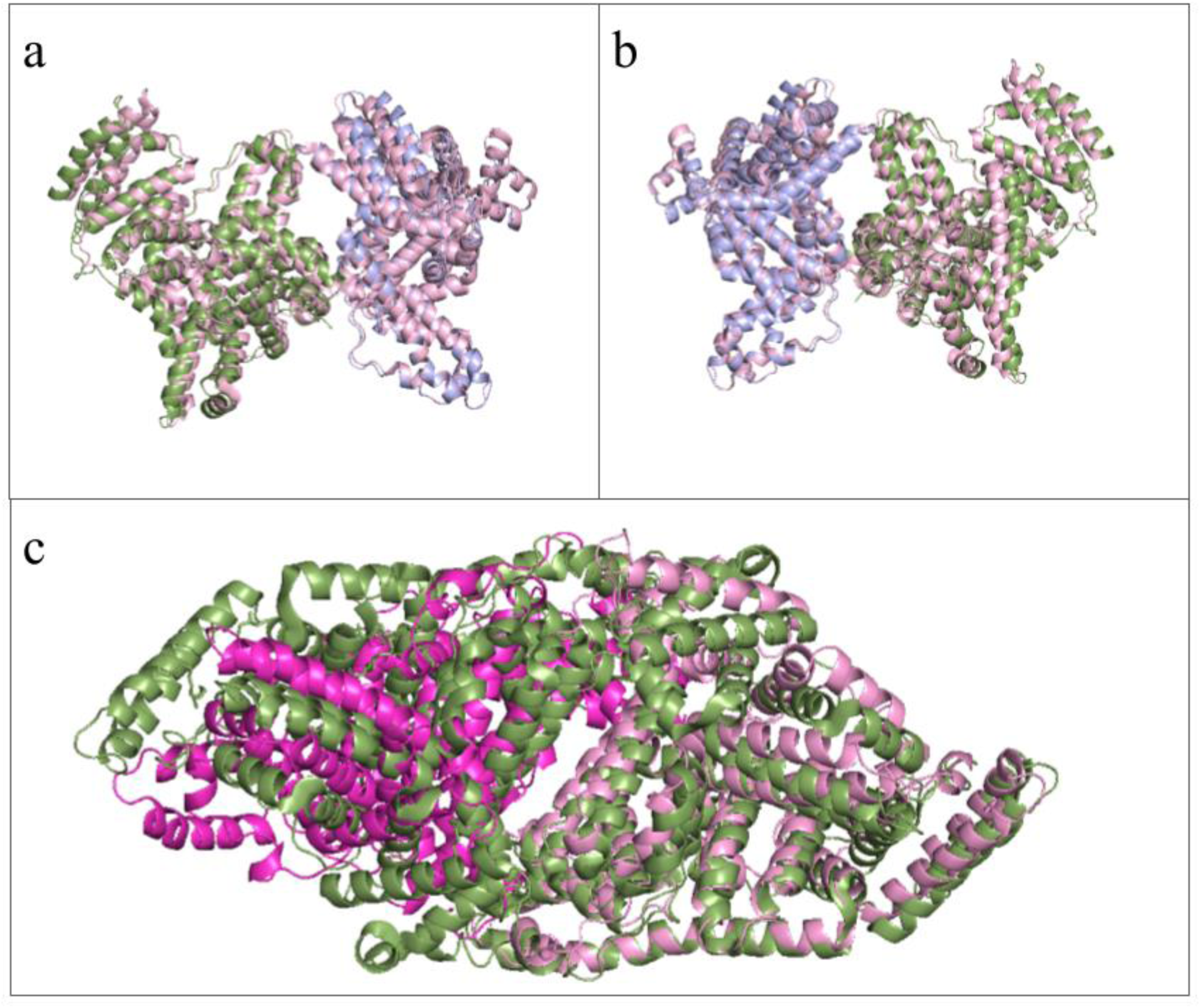
Our structure aligned to the other similar interface area structures 5Z0B (RMSD 1.0013) and 8CKS (RMSD 2.976), (a) the solved structure’s A (blue) and B (green) chains aligned to 5Z0B (pink) dimer, front view. (b) the solved structure’s A (blue) and B (green) chains aligned to 5Z0B dimer, back view. (c) the solved structure’s C1 (pink) and C2 (magenta) chains aligned to 8CKS (green) dimer, front view.

**Table 1:** The PISA-proposed salt bridges between both A-B chains and two C chains listed with bonding atoms and bond distances present.

|  | Chain B | Chain A | Distance (Å) |  | Chain C1 | Chain C2 | Distance (Å) |
| --- | --- | --- | --- | --- | --- | --- | --- |
| 1 | LYS 212<br>[ NZ ] | GLU 208<br>[ OE1 ] | 3.16 | 1 | LYS 93<br>[ NZ ] | ASP 308<br>[ OD1 ] | 3.49 |
| 2 | LYS 212<br>[ NZ ] | GLU 208<br>[ OE2 ] | 3.99 | 2 | LYS 205<br>[ NZ ] | GLU 321<br>[ OE2 ] | 3.86 |
| 3 | GLU 208<br>[ OE1 ] | LYS 212<br>[ NZ ] | 2.63 | 3 | LYS 212<br>[ NZ ] | GLU 208<br>[ OE1 ] | 3.58 |
| 4 | GLU 208<br>[ OE2 ] | LYS 212<br>[ NZ ] | 3.71 | 4 | ASP 308<br>[ OD1 ] | LYS 93<br>[ NZ ] | 3.49 |
|  |  |  |  | 5 | GLU 321<br>[ OE2 ] | LYS 205<br>[ NZ ] | 3.86 |
|  |  |  |  | 6 | GLU 208<br>[ OE1 ] | LYS 212<br>[ NZ ] | 3.58 |

**Table 2:**
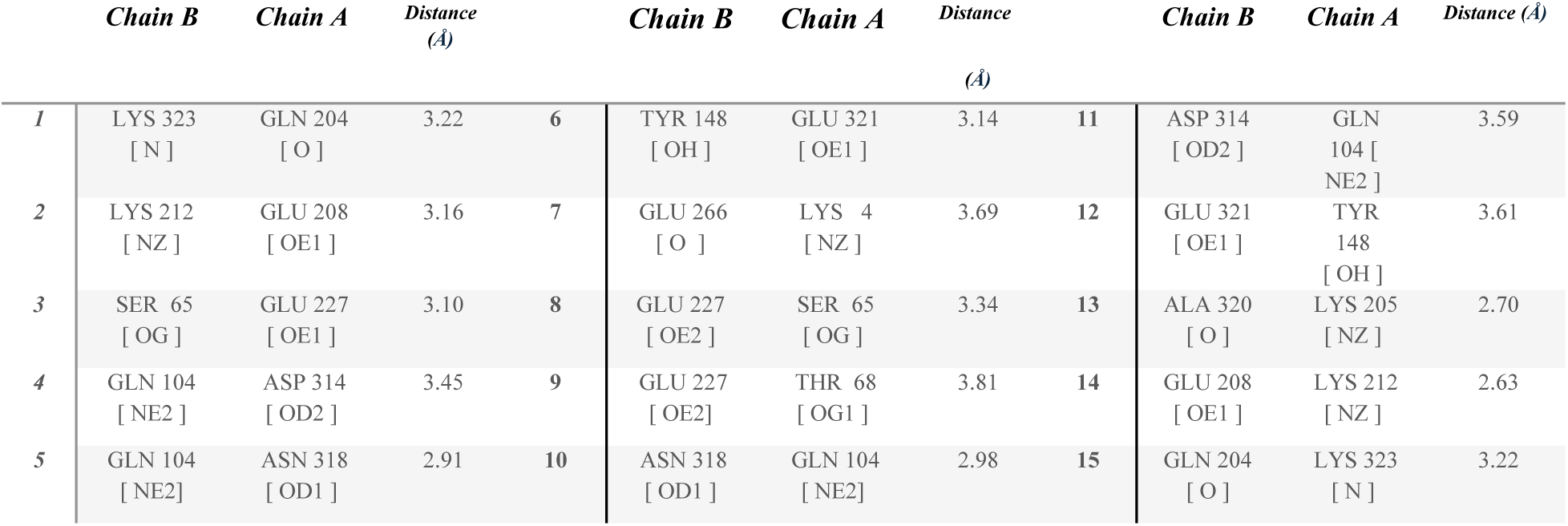
The PISA-proposed hydrogen bonds between A and B chains listed with bonding atoms and bond distances present.

**Table 3:**
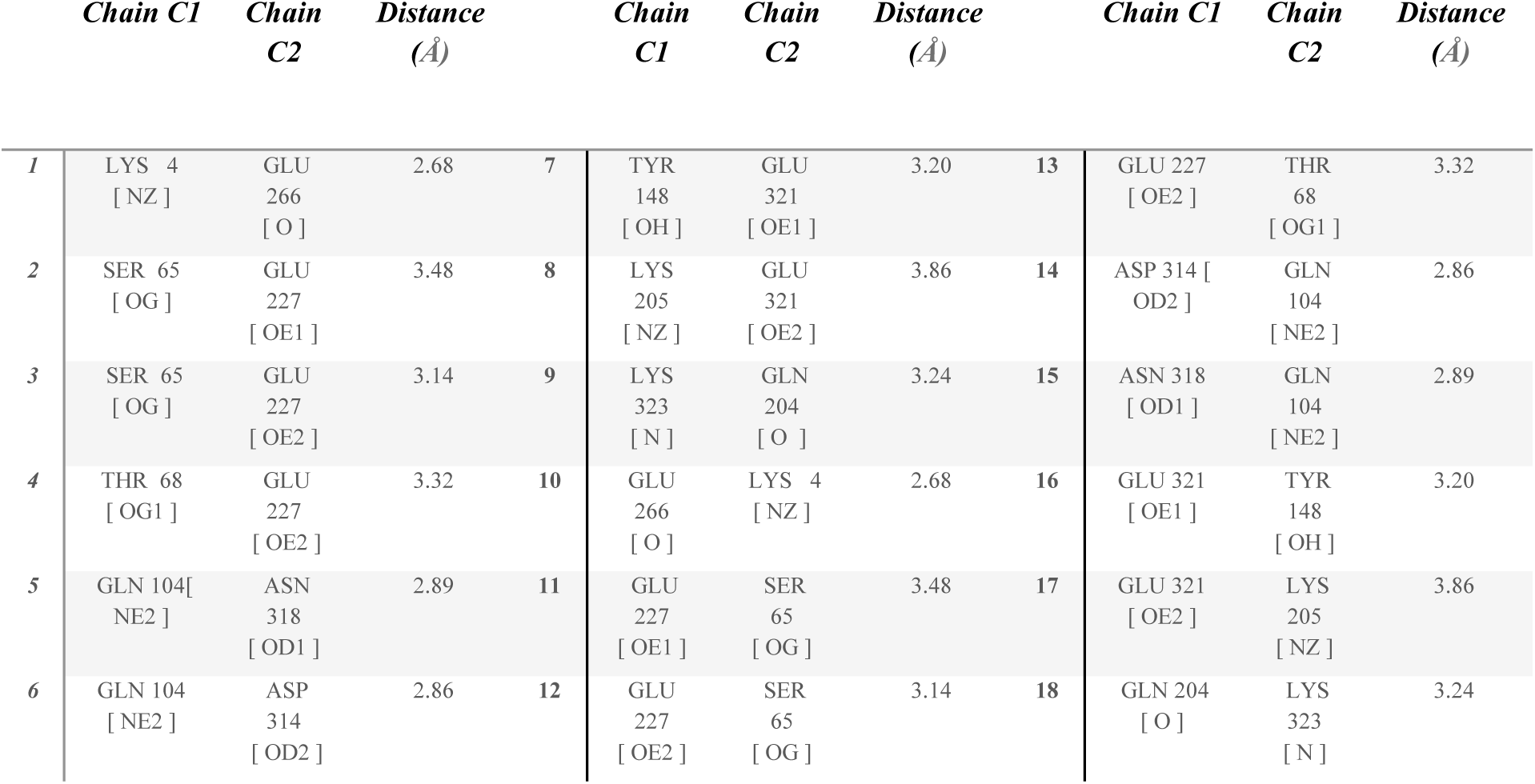
The PISA-proposed hydrogen bonds between the two C chains listed with bonding atoms and bond distances presented.

### Molecular Docking

The crystallized ligands warfarin, indoxyl sulfate, and 9-amino-camptothecin were docked into their corresponding active sites. After the docking procedure, the positions of identical atoms between the crystallized ligand and the docked ligand were compared, and the parameter of RMSD was calculated. RMSD calculation is a crucial method for validating the docking procedure [32], and an RMSD value under 2Å indicates that the docking procedure was close enough to the experimental result. Using the DockRMSD script [33], the RMSD value of our docked ligand was calculated. The calculated RMSD values are given in Table 4.

**Table 4.**
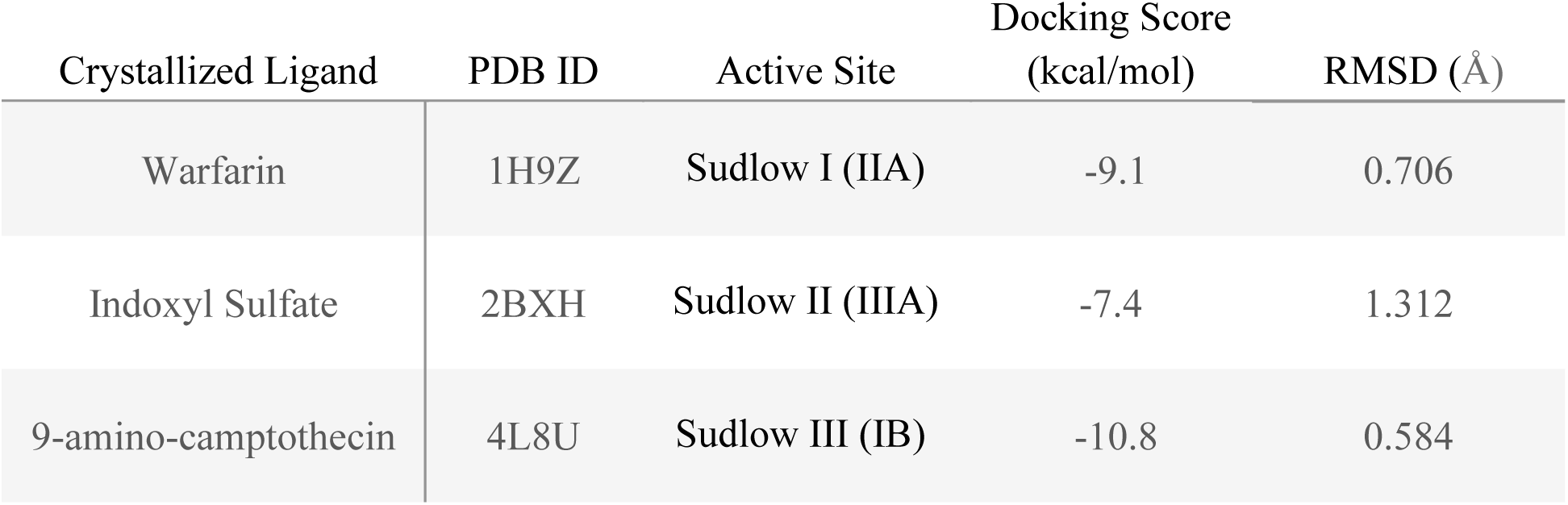
Docking results of crystallized ligands and RMSD values.

| Crystallized Ligand | PDB ID | Active Site | Docking Score<br>(kcal/mol) | RMSD (Å) |
| --- | --- | --- | --- | --- |
| Warfarin | 1H9Z | Sudlow I (IIA) | -9.1 | 0.706 |
| Indoxyl Sulfate | 2BXH | Sudlow II (IIIA) | -7.4 | 1.312 |
| 9-amino-camptothecin | 4L8U | Sudlow III (IB) | -10.8 | 0.584 |

After the validation of the docking procedure, dipyridamole is docked into different binding sites of HSA. The binding energies of dipyridamole to the Sudlow I, Sudlow II, and Sudlow III sites are given in the following Table 5.

**Table 5.** Docking scores of dipyridamole in different active sites.

| PDB ID | Active Site | Docking Score (kcal/mol) |
| --- | --- | --- |
| 1H9Z | Sudlow I (IIA) | -7.4 |
| 2BXH | Sudlow II (IIIA) | +10.4 |
| 4L8U | Sudlow III (IB) | -7.6 |

Molecular docking results show that dipyridamole demonstrates significant affinity towards the Sudlow Site I and Sudlow Site III of the HSA protein by forming several intermolecular interactions with the residues of the active sites. Furthermore, it has been shown that dipyridamole has no affinity towards the Sudlow Site II due to high intermolecular repulsions and the small size of the binding site.

### Sudlow Site I

Molecular Docking of dipyridamole towards the Sudlow Site I in the IIA subdomain of HSA protein has shown that dipyridamole has demonstrated significant binding affinity to the Sudlow Site I, as shown in Figure 16. The 2D map of dipyridamole and Sudlow Site I shows that dipyridamole’s aromatic rings form strong π-π interactions with the HIS242 residue and π-alkyl interactions between TRP 214, ALA215, LEU219, LEU238, and ALA291 residues. Furthermore, dipyridamole forms strong hydrogen bonds with LYS195, ARG257, and ASP451 residues with its hydroxyl tails at the two ends of dipyridamole.

**Figure 16:**
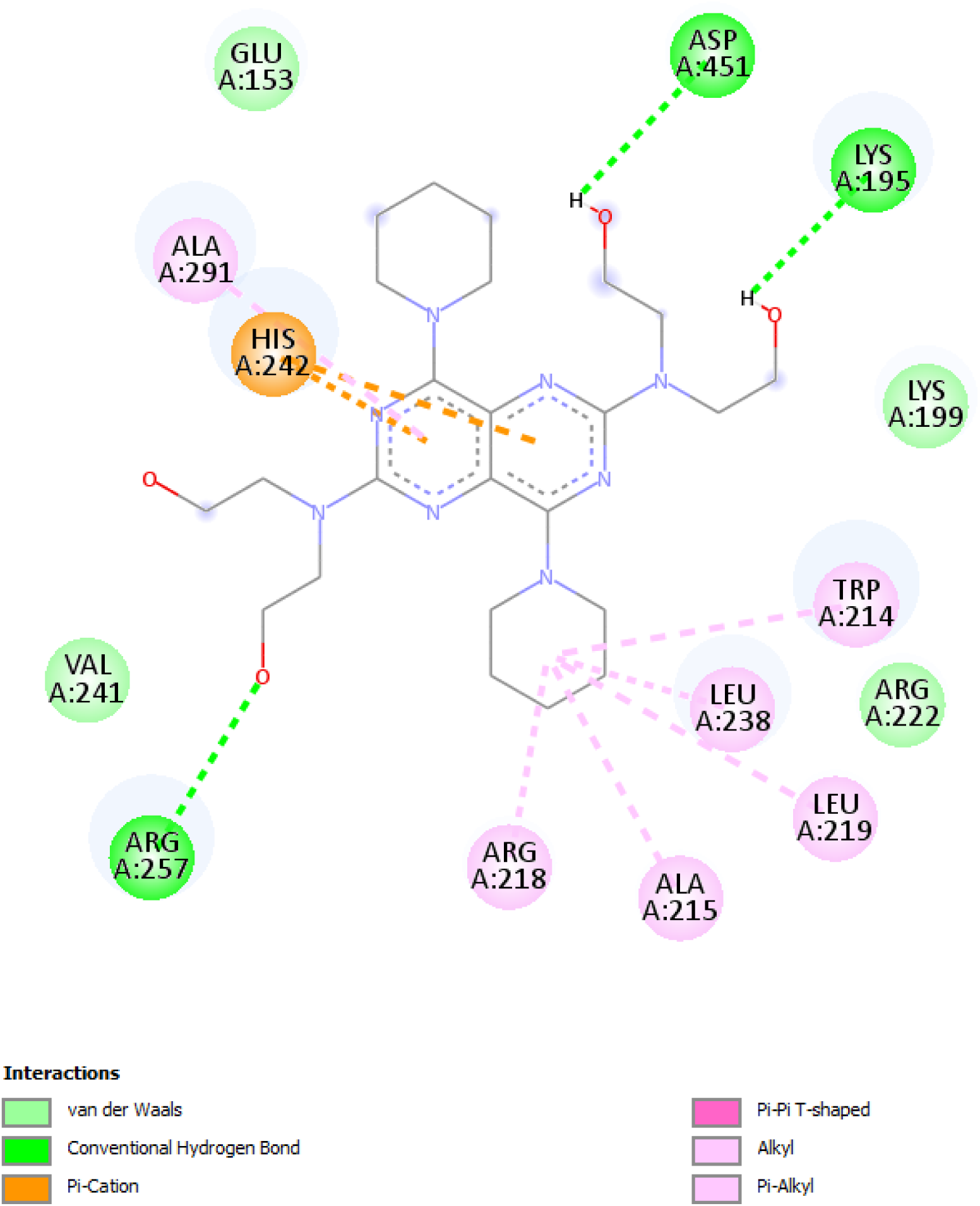
The 2D interaction map between dipyridamole and the amino acid residues of the Sudlow Site I is depicted.

**Figure 17:**
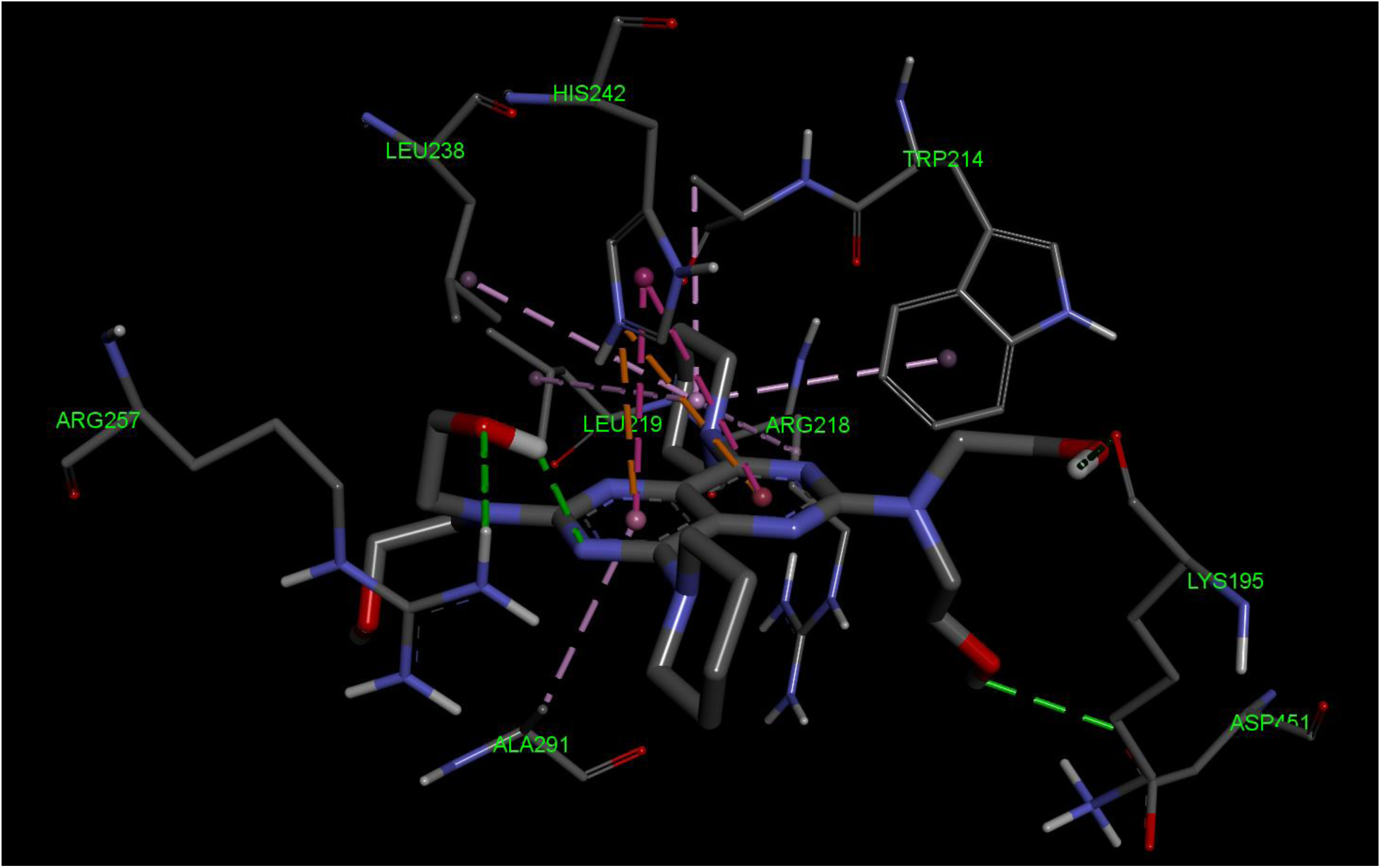
The 3D interaction map between dipyridamole and the amino acid residues of the Sudlow Site I is depicted.

### Sudlow Site III

Molecular docking of dipyridamole towards the Sudlow Site III in the IB domain of HSA protein shows that dipyridamole demonstrates significant affinity towards the Sudlow Site III, as shown in Figure 18. As depicted in the 2D interaction map, dipyridamole forms a strong hydrogen bond between its hydroxy tail and the ARG117 residue of the active site. Furthermore, dipyridamole forms several π interactions between its aromatic ring and heteroatomic saturated ring and the LEU115, HIS146, TYR161, ILE142, LEU182, LEU185, LEU190 residues of the active site.

**Figure 18:**
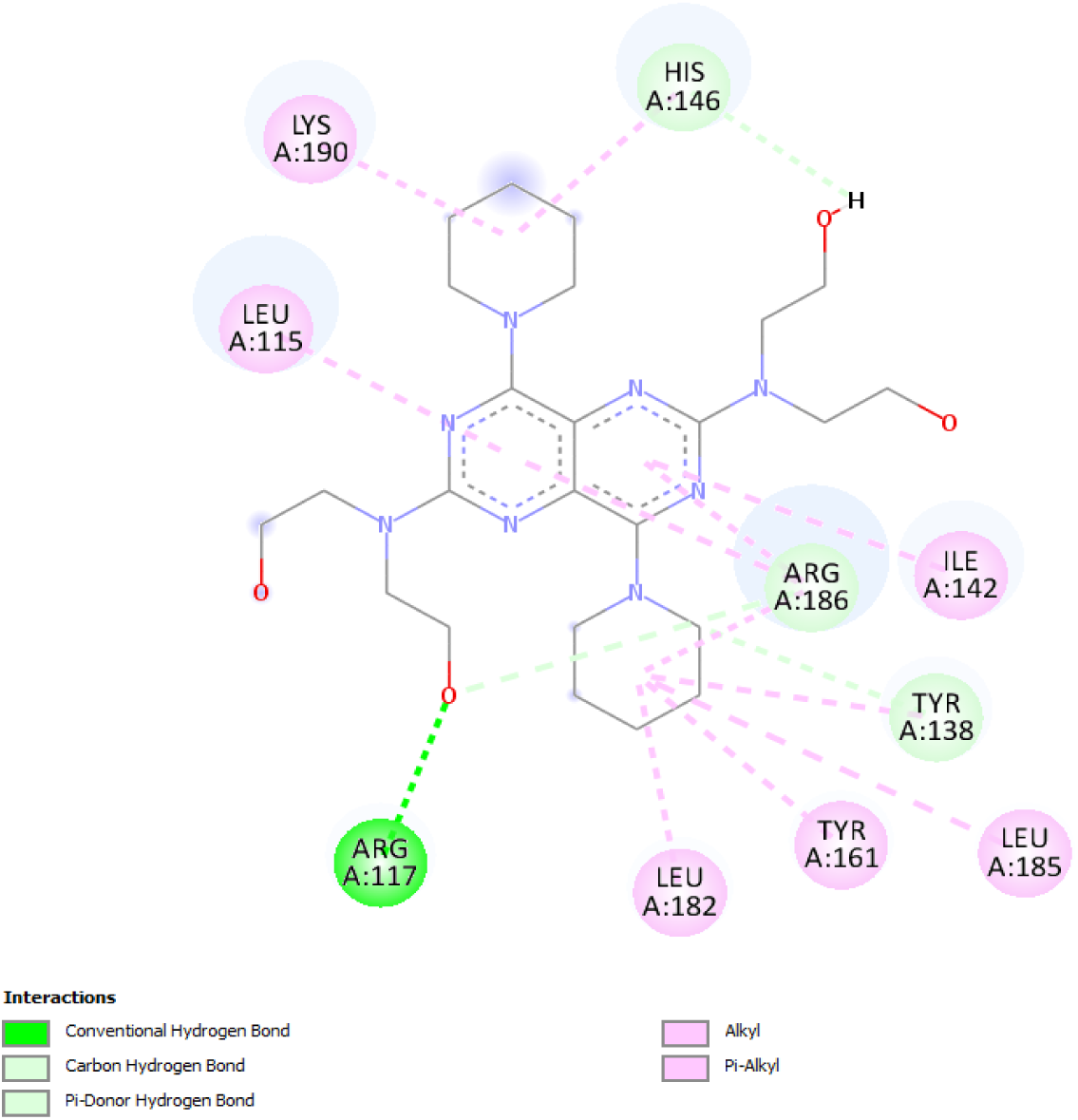
The 2D interaction map between dipyridamole and the amino acid residues of the Sudlow Site III is depicted.

**Figure 19:**
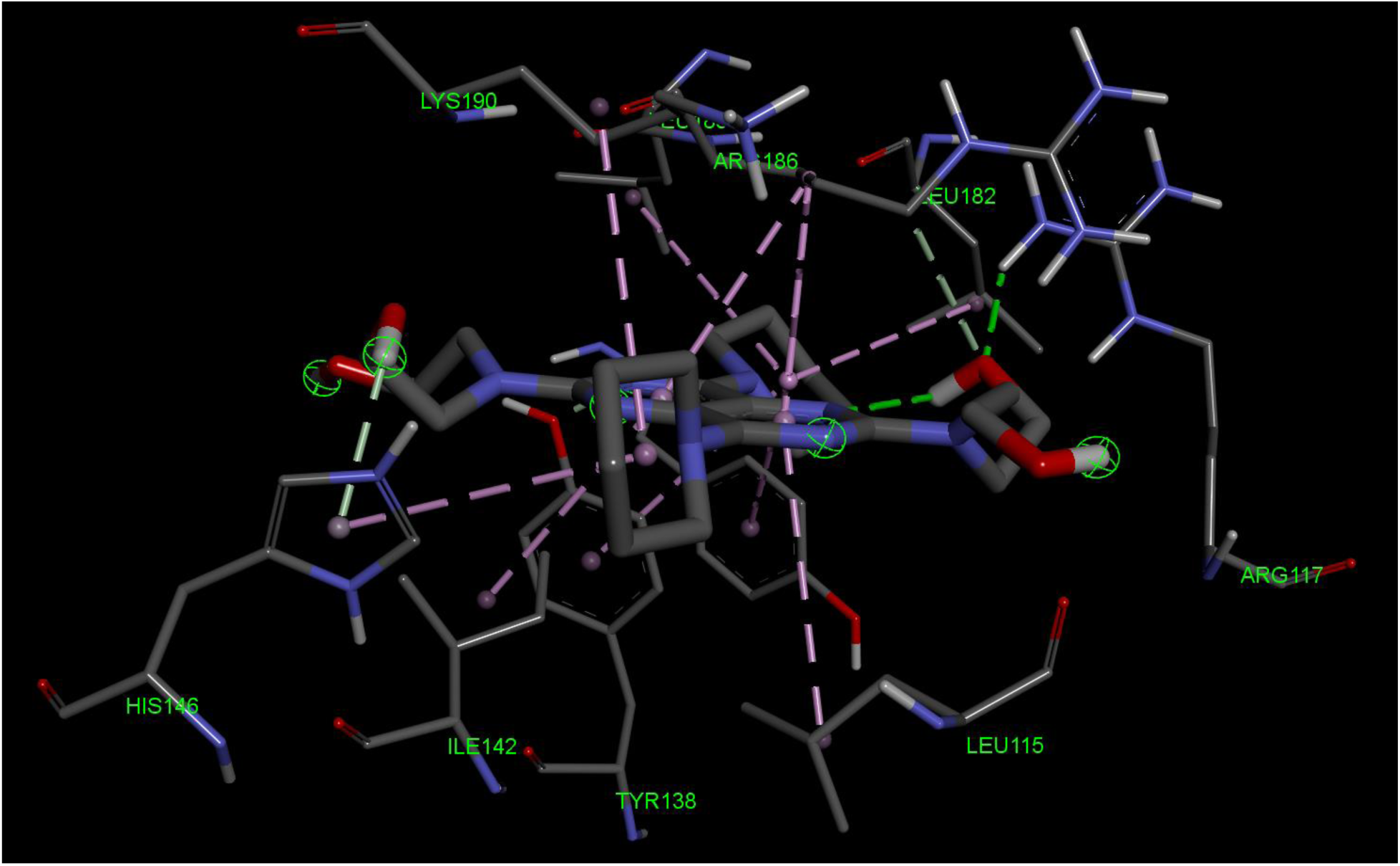
The 2D interaction map between dipyridamole and the amino acid residues of the Sudlow Site III is depicted.

## 4. Discussion

HSA dimerization patterns can offer novel insights into nanoparticle generation and recombinant-HSA-dependent delivery systems hence this research demonstrates two particular patterns of dimerization. Protein dimerization can be facilitated by electrostatic interactions hence the salt bridges and hydrogen bonds in the proposed HSA dimerization are characterized and identified as seen in Table 1, Table 2, and Table 3 [34].

The interface area histogram in Figure 8 shows that the distribution of the area in HSA crystals is arranged in two distinct normal distribution peaks the first one consisting of monomer crystallization and the second smaller one suggesting dimerization occurring before crystallization nucleation, the two peaks having a large gap between suggests the events leading to crystallization are different. The concentration of the unusual space groups such as *I 41* and *I 1 2 1* is also a notable difference further suggesting the crystallized units are different.

The overwhelming interface area of 9V61 structure and comparatively large number of electrostatic interactions suggest [35] an inherent interaction in the dimers that is different from crystallization artifacts suggesting somewhere in the process HSA monomer-dimer equilibrium shifted to favor dimers more. Once compared with the other structures with unusually high interface area in Figure 15, it is observed that there are differences between the structures even though interacting surfaces are the same with slight interacting residue differences between them. In Figures 11-14, the electrostatic interactions were given with the electron clouds visible to ensure this protein folding is confident in the interacting residues and is observed and not calculated during the molecular replacement.

In HSA-based drug delivery, it is important to tailor the approach based on the drug and the drug’s binding site on the protein, our dimerization patterns do not alter the sudlow drug binding sites and hence would not negatively affect the bindings of most drug-active ingredients. It may also be possible to alter the strength, association, and dissociation patterns of HSA nanoparticles and recombinant-dimer-HSAs via altering the identified residue using mutagenesis or targeted post-translational modifications. The increasing knowledge of possible HSA dimerization patterns may introduce new parameters, the protein itself, to the area of HSA nanoparticles hence we recommend nanoparticle efficiency comparison research with HSA mutants may lead to important steps in the field.

Molecular docking results show that dipyridamole shows significant affinity towards the Sudlow site I and Sudlow site III binding regions, whilst showing poor affinity towards the Sudlow site II. Dipyridamole’s low affinity towards the Sudlow Site II might be due to the small size of the Sudlow Site II, resulting in an unfavorable energy calculation in the docking process. Furthermore, it can also be deduced that binding events may affect the distribution of dipyridamole in the bloodstream, resulting in a direct effect on the pharmacokinetic profile of dipyridamole [36].

## 5. Conclusion

The identification and characterization of two novel HSA dimerization interfaces of 9V61 provide compelling evidence for previously unregarded plasticity in albumin self-interaction. The extensive electrostatic and hydrogen bonding interactions, coupled with their large interface surface areas, suggest that these dimers may hold biological or application-relevant significance rather than mere crystallographic artifacts. As the interactions listed in Table 1, Table 2, and Table 3 do not compromise known pharmacologically relevant binding domains, they present promising frameworks for engineering stable HSA-based nanoparticles. Moreover, these dimerization patterns propose a new variable in the field of albumin nanoparticle engineering—namely, the modulation of protein-protein interfaces. Future investigations into mutagenesis or post-translational modifications targeting the residues involved in these dimer interfaces may yield tunable nanoparticle constructs with altered stability and targeting efficiency. These insights not only contribute to the structural biology of HSA but also reinforce the usage of HSA dimers in advanced drug delivery platforms. As it comes to the dipyridamole interaction with Sudlow’s site I and Sudlow site III, we have calculated comparative binding energies between priorly established ligands indicating dipyridamole is expected to bind both Sudlow’s site I and Sudlow’s site III.

## Notes

### Competing Interest Statement

The authors have declared no competing interest.

